# Probing mechanical selection in diverse eukaryotic genomes through accurate prediction of 3D DNA mechanics

**DOI:** 10.1101/2024.12.22.629997

**Authors:** Jonghan Park, Galina Prokopchuk, Andrew R. Popchock, Jingzhou Hao, Ting-Wei Liao, Sophia Yan, Dylan J. Hedman, Joshua D. Larson, Brandon K. Walther, Nicole A. Becker, Aakash Basu, L. James Maher, Richard J. Wheeler, Charles L. Asbury, Sue Biggins, Julius Lukeš, Taekjip Ha

## Abstract

Connections between the mechanical properties of DNA and biological functions have been speculative due to the lack of methods to measure or predict DNA mechanics at scale. Recently, a proxy for DNA mechanics, cyclizability, was measured by loop-seq and enabled genome-scale investigation of DNA mechanics. Here, we use this dataset to build a computational model predicting bias-corrected intrinsic cyclizability, with near-perfect accuracy, solely based on DNA sequence. Further, the model predicts intrinsic bending direction in 3D space. Using this tool, we aimed to probe mechanical selection - that is, the evolutionary selection of DNA sequence based on its mechanical properties - in diverse circumstances. First, we found that the intrinsic bend direction of DNA sequences correlated with the observed bending in known protein-DNA complex structures, suggesting that many proteins co-evolved with their DNA partners to capture DNA in its intrinsically preferred bent conformation. We then applied our model to large-scale yeast population genetics data and showed that centromere DNA element II, whose consensus sequence is unknown, leaving its sequence-specific role unclear, is under mechanical selection to increase the stability of inner-kinetochore structure and to facilitate centromeric histone recruitment. Finally, *in silico* evolution under strong mechanical selection discovered hallucinated sequences with cyclizability values so extreme that they required experimental validation, yet, found in nature in the densely packed mitochondrial(mt) DNA of *Namystynia karyoxenos*, an ocean-dwelling protist with extreme mitochondrial gene fragmentation. The need to transmit an extraordinarily large amount of mtDNA, estimated to be > 600 Mb, in combination with the absence of mtDNA compaction proteins may have pushed mechanical selection to the extreme. Similarly extreme DNA mechanics are observed in bird microchromosomes, although the functional consequence is not yet clear. The discovery of eccentric DNA mechanics in unrelated unicellular and multicellular eukaryotes suggests that we can predict extreme natural biology which can arise through strong selection. Our methods offer a way to study the biological functions of DNA mechanics in any genome and to engineer DNA sequences with desired mechanical properties.

---

Present-day genomes are layered with multiple ‘codes’, including the genetic code in protein-coding regions, transcription factor motifs in regulatory regions, genomic code for nucleosome positioning and a histone code. Might there be another layer of ‘mechanical code’ where specific DNA sequences and chemical modifications directly influence biological functions through DNA mechanical properties^1,2^? Previously, we extended the DNA cyclization assay, first developed in 1981^3^, to single-molecule resolution and reported a profound effect of sequence on cyclization rate, which is a measurable proxy for DNA mechanics such as anisotropic bending propensity and intrinsic flexibility^4^. Subsequently, loop-seq, a sequencing-based readout of single DNA cyclization, enabled genome-scale quantification of intrinsic cyclizability^5^ and revealed highly rigid DNA near gene promoters and a rigidifying effect of cytosine methylation^1,5^. Based on loop-seq data, bioinformatics models have been developed to predict cyclizability from sequence and infer DNA mechanics across key genomic landmarks in diverse genomes^6–9^. But as we will show below, the existing models have limited accuracy, did not correct for biases introduced during loop-seq, and did not consider spatial anisotropy in DNA bending.

Here, we developed an approach to predict bias-corrected DNA cyclizability with > 95% accuracy and to infer intrinsic bending direction from sequence alone. We demonstrate the power of the approach in three novel applications: (1) spatial analysis to predict intrinsic bending direction, (2) probing mechanical selection from population genetics, and (3) discovery of extreme naturally occurring DNA mechanics from its *in silico* evolution.

## Accurate cyclizability prediction

For reliable prediction of the mechanical properties of DNA of any sequence, we need a large dataset linking DNA sequence to its mechanics. A proxy for DNA mechanics, intrinsic cyclizability, abbreviated as C0, was previously determined using loop-seq (Fig. 1a, Supplementary Fig. 1)^5^. C0 values of several libraries containing more than 150,000 50-bp sequences, randomly generated or derived from the yeast genome, are currently the standards against which the performance of sequence-to-cyclizability prediction is evaluated^6–9^. Because low sequencing read counts reduced the accuracy of some C0 measurements (Supplementary Fig. 2), we removed C0 values of high uncertainty caused by low read counts (Methods, Supplementary Fig. 3). Using the refined dataset, our deep-learning approach achieved a Pearson’s correlation of ∼0.96 between measured and predicted C0 (Fig. 1b, Supplementary Fig. 4), much above the previous analyses whose maxima were 0.75∼0.77 (Fig. 1c).

**Fig. 1.**
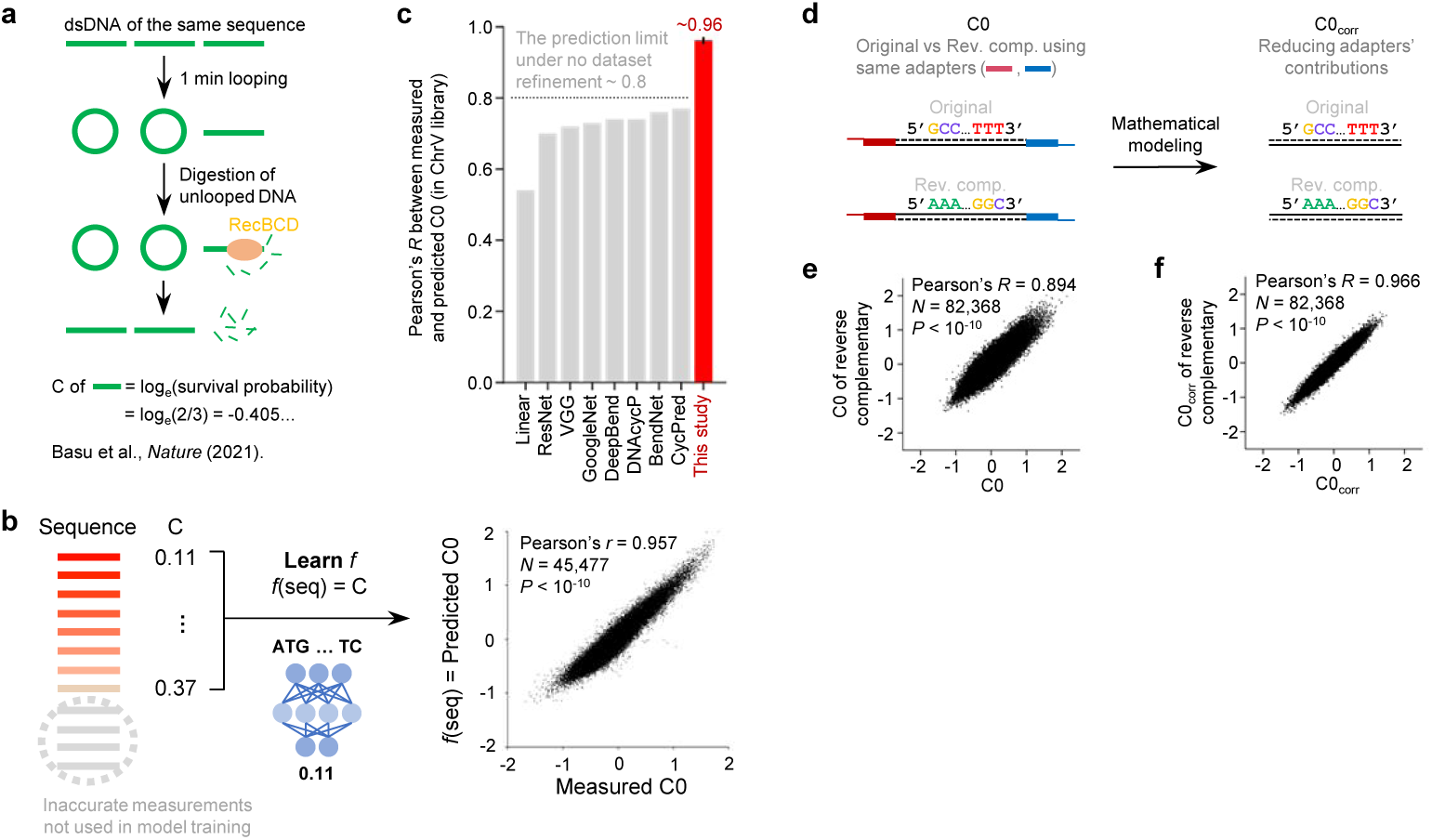
Highly accurate prediction of cyclizability. **a**, Schematic of loop-seq and cyclizability measurement (adopted from Basu et al^5^). **b**, Schematic of the model training process and scatter plot showing measured vs predicted C0 for the training dataset from the Tiling library^5^. A detailed schematic is found in Supplementary Fig. 3b. For model performance on the testing datasets, see Supplementary Fig. 4a. Pearson’s correlation, sample size, and the corresponding two-tailed *P*-value are shown. **c**, Performance of our prediction on the ChrV dataset (red) relative to previous predictions^1,6–9^ (gray). Error bar is a 95% confidence interval of Pearson’s correlation coefficient *R*. **d**, Schematic of adapter-dependent (C0) and adapter-corrected (C0_corr_) cyclizability. **e**, C0 values of the original vs the reverse complementary sequence of Tiling library. **f**, C0_corr_ of the original vs the reverse complementary sequence of Tiling library. **e**, **f**, Pearson’s correlation coefficient *R*, sample size, and the related two-tailed *P*-value are shown.

Loop-seq requires a pair of adapter sequences attached to the ends of the variable sequences (50 bp in most studies, Fig. 1d, Supplementary Fig. 1). The adapters can create bias by bending synergistically with the variable sequence. Indeed, C0 shows relatively weak correlation between the original sequences and their reverse complement (Fig. 1e)^5^, likely because the swapped adapter sequences impose a different mechanical context. Therefore, we devised a mathematical correction to remove the contribution of the adapter sequences (Methods, Supplementary Fig. 5). This improved the Pearson’s correlation of the cyclizability between the original sequence and its reverse complement to 0.97 from 0.89 (Fig. 1e, f). We termed C0 after the correction ‘adapter-corrected intrinsic cyclizability’, C0_corr_, which should be a quantity independent of its mechanical context. More specifically, C0_corr_ is independent of sequence orientation (Fig. 1f) and rotational phasing (Supplementary Fig. 5c). Thus, we will henceforth use ‘cyclizability’ to refer to adapter-corrected intrinsic cyclizability (C0_corr_). Lastly, we developed heuristic algorithms to enhance the speed of cyclizability predictions across genomic datasets spanning multiple Gbps (Supplementary Note 8).

### Spatial analysis reveals intrinsic bending direction

Because loop-seq was performed with DNA tethered to a bead surface, for sequences that prefer to bend toward the tethering point, steric hindrance will lower their apparent cyclizability (Fig. 2a)^5^. In a previous study, loop-seq analyses with three different tether positions were implemented to account for this effect and yielded intrinsic cyclizability C0, which is independent of tether position^5^. Using the same dataset but after correcting for the adapter effects, we developed a ‘spatial analysis’ which allowed us to predict the preferred direction of bending at every position on the DNA (Methods). As an example, for the well-positioned nucleosomes in budding yeast (*Saccharomyces cerevisiae*) *in vivo*^10^, we found that the preferred bending direction of their wrapped DNA is towards the center of the nucleosome (Fig. 2b, see red lines overlaid on nucleosome structure), suggesting that nucleosomes in yeast favors genomic DNA with static bending directions that are compatible with the sharply bent DNA conformations found in nucleosome structures^11^.

**Fig. 2.**
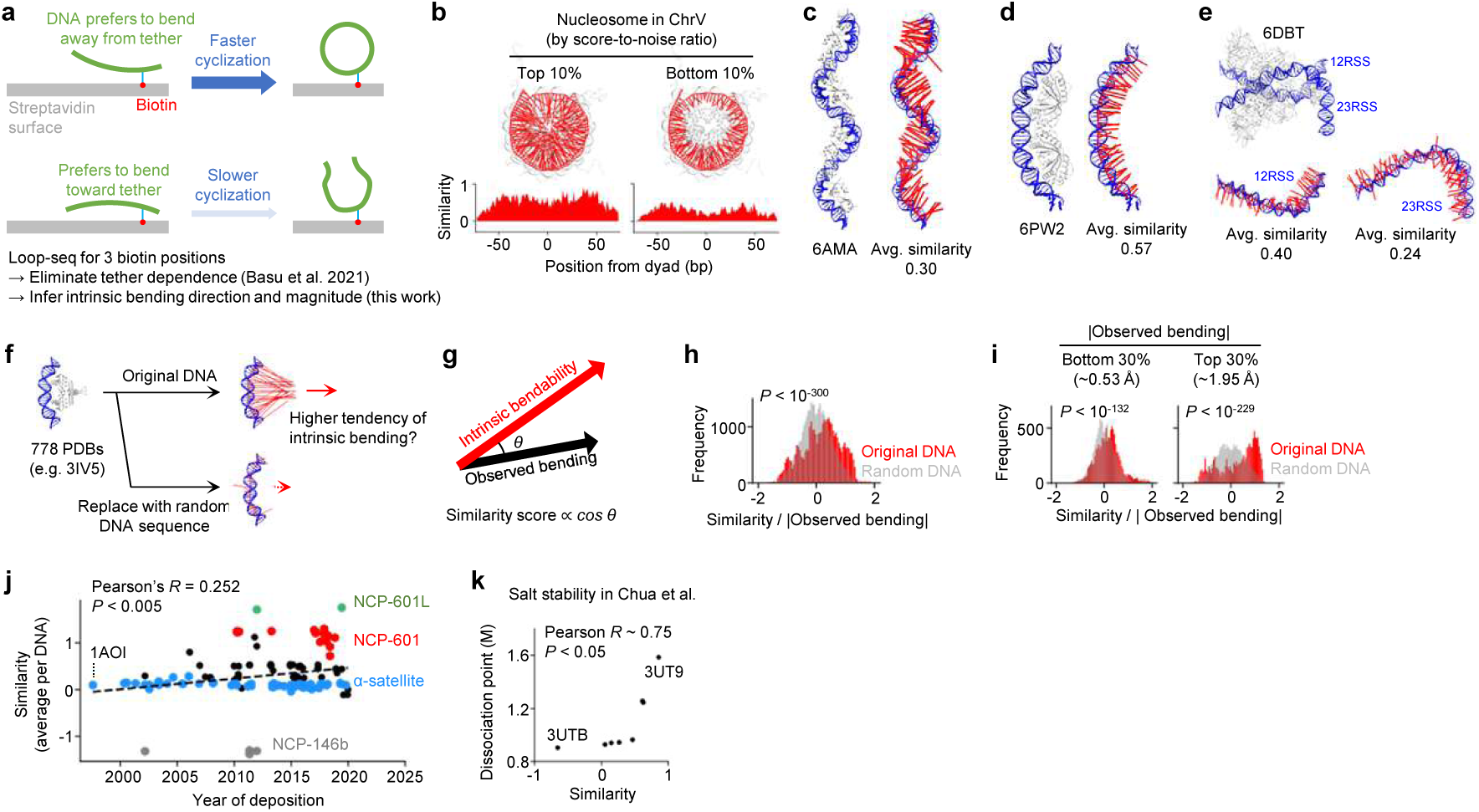
Selection of DNA mechanical properties in available structures. **a**, Cyclization rate is affected by anisotropic DNA bending in space. **b**, DNA bending of yeast nucleosome DNA with top or bottom 10% score-to-noise ratio for the dyad positions inferred from chemical mapping data^10,77^. The magnitude of predicted bending as denoted as red lines was scaled by a factor of 100 when visualized in 3- dimensional space (Methods). **c**, **d**, **e**, 3-dimensional DNA bending of protein-DNA complexes overlaid with the original structures. PDB IDs 6AMA, 6PW2, and 6DBT were used, respectively. **f**, Bending of original DNA sequences used in the reported structures in protein data bank (PDB) vs randomized sequences in 3-dimensional space. The predicted direction of DNA bending is shown in red, and the line length is proportional to the magnitude of bending predicted (Methods). **g**, Similarity score is an inner product of vectors for intrinsic bending and observed bending (Methods). Selection of DNA mechanical properties for all (**h**), as well as for the most bent (top 10%) and the least bent (bottom 10%) DNA molecules (**i**), within the curated set of protein-DNA structures in RCSB database. Mechanical properties were evaluated using a normalized similarity score, which is similarity divided by DNA bending vector length (Methods). **j**, Similarity score averaged over each DNA in the published nucleosome structures vs publication year. Black symbols are for nucleosome structures whose sequence could not be assigned to one of the annotated sequences denoted by colored symbols. **k**, Similarity score averaged over each nucleosome DNA vs salt stability measured in Chua et al^16^.

Is it possible that highly bent DNA molecules in known protein-DNA structures intrinsically prefer to bend in the same direction? A spectacular affirmative example is the BldC protein of *Streptomyces* that bends DNA spirally (Fig. 2c)^12^. Intrinsic bending direction overlaid on the structure matches the observed DNA bending. A second example is EBNA1 of Epstein-Barr virus that bends DNA in the direction of its intrinsic bending propensity (Fig. 2d)^13^. A third example is RAG1/2 recombinase where the deformation of 23 bp recombination signal sequence for V(D)J recombination agrees with intrinsic DNA bending (Fig. 2e)^14^. We developed a convenient web interface for users to enter the PDB ID of a structure and interactively visualize intrinsic bending directions overlaid with the structure (Methods).

For a systematic investigation, we performed spatial analysis for 778 structures that contain dsDNA molecules with well-defined DNA sequence in the RCSB database^15^ (Methods). We compared the predicted 3D mechanics of DNA sequences used in the structural studies with those of randomly generated DNA of the same length (Fig. 2f) by calculating a similarity score: the inner product between vectors for the intrinsic bending direction and the observed bending direction (Fig. 2g, Methods). A positive similarity score implies that the observed bending is favored by intrinsic DNA mechanics. We found that DNA sequences used for experimental structural determination have higher similarity scores compared to random DNA (Fig. 2h), especially when the observed structure has strong DNA bending (top 30%, *P* < 10^-229^, Fig. 2i, right), but the effect is still significant for structures with low DNA bending (bottom 30%, *P* < 10^-132^, Fig. 2i, left). Even when we divided the structures into nucleosomes (n=140), transcription factors (n=275), and all others (n=322)), the similarity score was higher for the DNA sequences used for structural determination compared to random DNA sequences for each group (Supplementary Fig. 6a). Therefore, available structures have a significant bias in favor of sequences with predicted intrinsic bending direction that matches the observed bending direction. Such biases were dependent on the structural category, largest for nucleosomes (*P* < 10^-167^), smallest for transcription factors (*P* < 10^-17^), and the remainder intermediate (*P* < 10^-74^) among the top 30% most bent DNA structures (Supplementary Fig. 6a). Overall, our spatial analysis applied to the available structures suggests that many proteins co-evolved with their DNA partners to recognize and bend DNA sequences according to their intrinsic bending direction.

Can some of the biases be due to ‘selection’ by the researchers, for example, due to increased stability of complexes that facilitates structural analysis? The answer seems to be yes for the nucleosome structures because the average similarity score for deposited nucleosome structures has increased over time (Fig. 2j). Indeed, there is a strong positive correlation between the similarity score and salt stability of nucleosomes^16^ (Fig. 2k). The first reported nucleosome structure used alpha-satellite DNA^11^ with negative similarity scores near the dyad (Supplementary Fig. 6b). In contrast, Widom 601 DNA^17^, widely adopted for more recent studies^18–20^, has positive similarity scores in almost all positions (Supplementary Fig. 6c). Higher similarity scores on the left side of Widom 601 DNA might explain the asymmetrical unwrapping of nucleosome DNA under tension^21^ (Supplementary Fig. 6c). Also, NCP-601L with uniformly high (Supplementary Fig. 6d) and NCP-146b with uniformly low similarity scores (Supplementary Fig. 6e) are also the nucleosomes with the highest and the lowest nucleosome stability against salt titration, respectively (Fig. 2k)^16^.

### Probing mechanical selection from population genetics

If certain mechanical properties are disfavored due to functional constraints, sequence variants responsible will rarely become fixed in natural populations. Borrowing from tools to quantify selection pressure on sequence variants in protein coding^22^ and non-protein coding elements^23,24^, we developed a method to quantify selection pressure acting on DNA mechanics from population genetics data (Fig. 3a, Methods) and applied it to the centromeric sequences of 1,011 isolates of *S. cerevisiae*^25^.

**Fig. 3.**
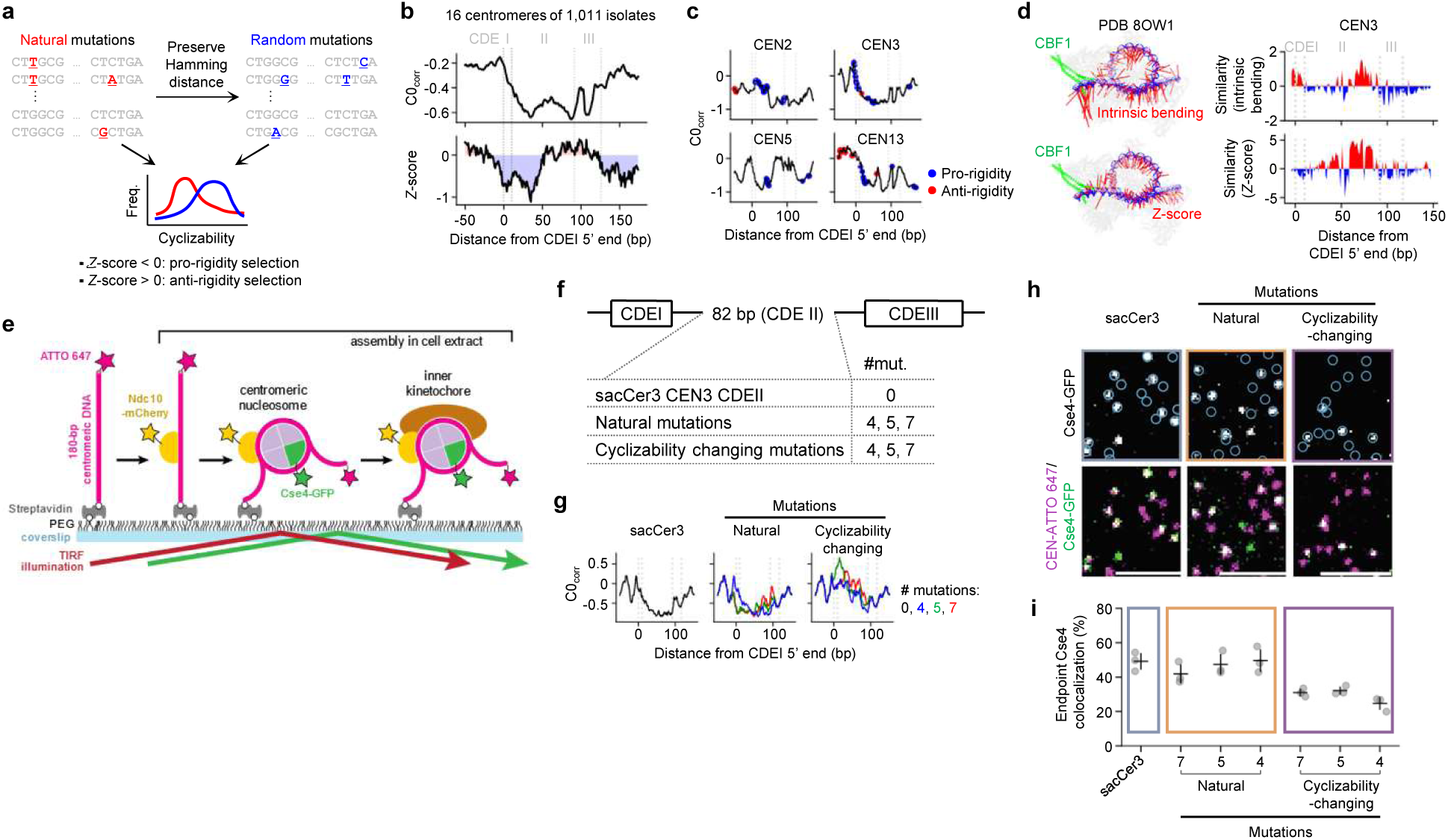
Mechanical selection in yeast centromeres modulates the stability of inner kinetochores. **a**, Schematic of the process quantifying mechanical selection using the populational genetics data of yeast^25^ (Methods). **b**, Cyclizability and mechanical selection (*Z*-score) averaged over 16 yeast centromeres collected from 1,011 yeast isolates. Regions of negative Z-score are highlighted blue. **c**, Mechanical selection in CEN2, 3, 5, and 13. Pro-and anti-rigidity selection are marked by red and blue colors, respectively (Methods). **d**, Intrinsic propensity of DNA bending (top) and mechanical selection (bottom) in inner kinetochore. The results are overlaid on the 3-dimensional structure of inner kinetochore (PDB 8OW1, left), and the corresponding similarity scores are plotted (right). **e**, Schematic of the fluorescent label location used in smTIRF colocalization assay. **f**, Overview of sacCer3, natural, and cyclizability-changing CDEII DNA mutants used for single molecule fluorescence colocalization analysis. **g**, Cyclizability of CDEII sequences in **f**. **h**, Example images of total internal reflection fluorescent microscopy endpoint colocalization analysis of visualized Cse4-GFP on sacCer3 CDEII DNA (left), CDEII with natural mutations (middle) or cyclizability-changing mutations (right), with colocalization shown in relation to identified DNA in blue circles. Bottom panels show respective overlays of DNA channel (magenta) with Cse4-GFP (green). Scale bars: 3 μm. **i**, Graph indicates quantification of endpoint colocalization of Cse4 on CDEII DNA (left), CDEII with natural mutations (middle) and cyclizability-changing mutations (left). Points indicate individual experiments (n=3) where ∼1,000 DNA molecules were identified per replicate.

After aligning the genome sequences of the 1,011 isolates at the 16 centromeres, we created, for each 50 bp window, a simulated sequence that matches the Hamming distance of the corresponding natural sequence from the consensus sequence (Fig. 3a, Methods). We compared cyclizability distributions to determine whether variations found in natural sequences deviate significantly from the simulated sequences. For example, significantly lower cyclizability in the natural sequences suggests that any mutation increasing cyclizability, thus reducing DNA rigidity, was deleterious, leading to ‘pro-rigidity selection’.

The centromere DNA is a site of kinetochore assembly which is critical for proper chromosome segregation^26–28^. Each chromosome in *S. cerevisiae* has a single centromere with three centromere defining elements (CDEI, CDEII, and CDEIII)^29,30^. CDEI and CDEIII contain the recognition sites of centromere binding factors^31–33^ but the consensus sequence for CDEII has not been identified^34^. CDEII has high content of polymeric runs of A or T^34^, and the longer A or T tracts were proposed to facilitate the deposition of centromeric nucleosome containing Cse4^CENP-A^ by an unknown mechanism^35^. We found that the centromeric DNA is rigid (average C0_corr_ ∼-0.5, Fig. 3b, top), possibly to prevent random nucleosomes from occupying the centromeres^36^. Averaged across all 16 centromeres, 1,011 yeast isolates showed pro-rigidity selection at CDEI and the upstream portion of CDEII (Fig. 3b, bottom), with a similar trend observed in individual centromeres (Fig. 3c).

As a further test, we examined CDEII bending in the inner-kinetochore structure^37^ by calculating the similarity score for the chromosome 3 centromere (CEN3) CDEII sequences of 1,011 yeast isolates. We found a positive similarity score in the region that curves around the centromeric nucleosome, which overlaps with the downstream portion of CDEII (50 ∼ 90 bp from the 5’ end of CDEI, Fig. 3d, top). The same region was under mechanical selection, accumulating variants with positive similarity scores (Fig. 3d, bottom, Methods). Therefore, for centromere function, intrinsic DNA mechanics in 3D, not just rigidity, appears to be under selection.

To probe the molecular basis underlying sequence-dependent centromere function, we measured the recruitment of the histone H3 centromeric variant Cse4 to CEN3 under a perturbation of DNA mechanics using single molecule fluorescence colocalization microscopy (Fig. 3e, Methods) ^35^. We prepared the wild type CEN3 and three natural mutants by selecting three variants among the natural population. The variants contained 4 to 7 mutations in CEN3 CDEII that preserve DNA mechanical properties. We also generated three mutants that contain the same number of mutations but do not preserve DNA mechanical properties (Fig. 3f, g, Supplementary Table 1). All three mechanics-preserving natural variants recruited Cse4 to a similar level as the wild type whereas all three cyclizability-changing mutants showed significantly reduced recruitment (Fig. 3h, i), supporting the importance of CDEII mechanics in Cse4 recruitment. As a control, an earlier step, the recruitment of Ndc10 to CDEIII, was unaffected by the change in CDEII mechanics (Supplementary Fig. 7a, b). Taken together, our mechanical selection analysis showed that the centromeric element CDEII has evolved to maintain specific mechanical properties important for kinetochore assembly.

### *In silico* evolution of DNA mechanics

To understand how desired mechanical properties may emerge from selection, we adopted the strong-selection weak-mutation approach^38^, previously used to link promoter sequence to gene expression in *S. cerevisiae*^24^. Starting from a 50 bp sequence chosen at random, all possible single substitutions were considered in each cycle to produce a total of 150 new sequences. The sequence with the highest and lowest predicted C0_corr_ value were chosen in cyclizability-maximizing or minimizing selection, respectively, while one of the 150 sequences was randomly picked to simulate genetic drift. The chosen sequence serves as an input for the subsequent cycle of simulation (Fig. 4a).

**Fig. 4.**
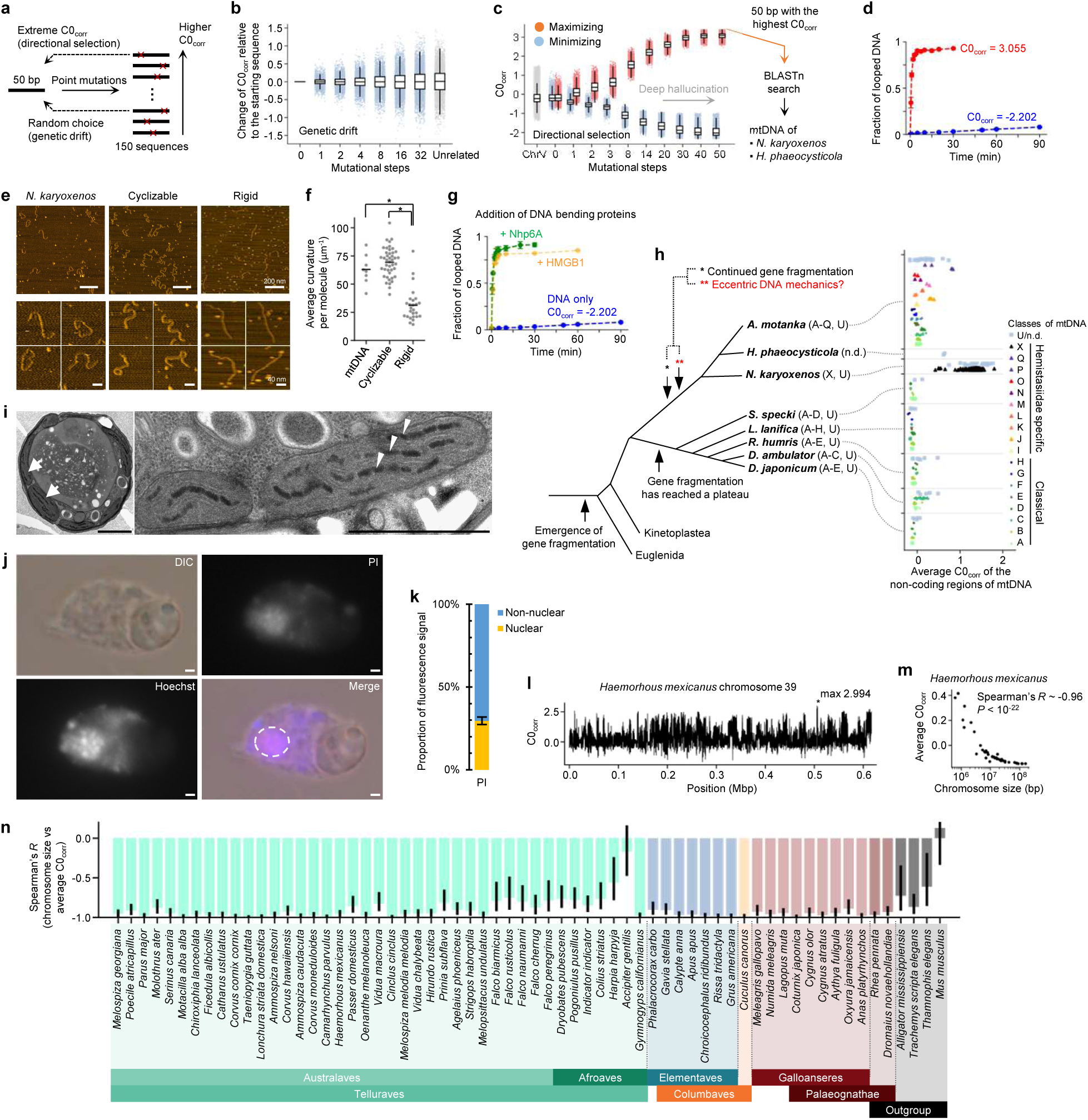
Evolution of extreme cyclizability in *Namystinia karyoxenos* mitochondrial DNA. **a**, Schematic of *in silico* evolution. Starting with 6,420 random 50 bp sequences, 150 point mutations are generated from each sequence. The mutation that finds the extrema of C0_corr_ (directional selection) or a random mutation (genetic drift) is selected as the input for the next round. **b**, Change in C0_corr_ relative to the starting sequence after 0, 1, 2, 4, 8, 16, and 32 mutational steps of genetic drift, starting with 6,420 random 50 bp sequences. Differences between the C0_corr_ of random 50 bp sequences and arbitrarily assigned starting sequences were plotted separately as an unrelated set. **c**, C0_corr_ after the indicated number of mutational steps of directional selection, starting from 6,420 random 50 bp sequences. Gray, C0_corr_ in yeast chromosome V, red, maximizing selection, blue, minimizing selection. **b**, **c**, Whisker box plots are shown together with the scattered data. **d**, The fraction of looped DNA molecules in the real-time single-molecule cyclization assay (Methods). 50 bp DNA sequences with extremely high (C0_corr_ ∼ 3.055) or low cyclizability (C0_corr_ ∼-2.202) were used. Error bars are standard deviations of three experiments. **e**, Example AFM images of a 600 bp linear segment of mitochondrial DNA of *N. karyoxenos* and two DNA molecules of the same length with extremely high or low cyclizability (Methods). Cyclizability vs position for the 600 bp DNA sequences is shown in Supplementary Fig. 9e. The top row shows the whole field (scale bar: 200 nm), and the bottom rows show zoom-ins (scale bar: 40 nm). **f**, Static curvature averaged per DNA molecule in AFM images (Methods). *P*-values (two-tailed t-test) lower than 0.05 are indicated as *. **g**, The fraction of looped DNA molecules in the real-time single-molecule cyclization assay after the addition of DNA-bending proteins (Methods). DNA with low cyclizability used in the previous cyclization assay in **d** was tested with the addition of Nhp6A (n=3 experiments) and HMGB1 (n=1 experiment). Error bars represent standard deviations of replicates. **h**, Cladogram depicting the phylogenetic relationships of diplonemid protists ^78^ and the average C0_corr_ in non-coding regions of mitochondrial genomes. Classes of mitochondrial genomes are noted next to species names. **i**, Transmission electron micrographs of *N. karyoxenos* displaying reticulated peripheral mitochondrial branches (arrows, left panel) and bead-like electron-dense mtDNA (arrowheads) located among the mitochondrial cristae (right panel). Scale bar: 1 µm. **j**, Light microscopy micrographs of *N. karyoxenos* labelled with the minor groove binder, Hoechst 33342, as well as with the base pair intercalating dye, PI. DIC - differential interference contrast. Scale bar: 1 µm. **k**, Proportion of nuclear and non-nuclear (mitochondrial and endosymbiont) DNA as measured from PI fluorescence signal. *n* = 372 cells, error bars represent standard error of proportion. **l**, Cyclizability along chromosome 39 of *H. mexicanus*. **m**, Average cyclizability vs length of each chromosome of *H. mexicanus*. Spearman’s *R* and the related two-tailed *P*-value are shown. **n**, Spearman’s *R* between average cyclizability vs length of each chromosome for 60 different bird species and four non-bird neighbors. Error bars indicate 95% CI.

In the genetic drift simulation, C0_corr_ diverged as mutations accumulated (Fig. 4b), and even two rounds of single point mutation were sufficient to change C0_corr_ by 0.77 in some sequences, which is nearly three times the standard deviation of the initial C0_corr_ distribution. In parallel, a series of maximizing and minimizing selections shifted the overall distributions of C0_corr_ toward the corresponding extrema (Fig. 4c). Thus, selection pressure can readily alter cyclizability with just a few mutations.

Continued directional selection until the 50^th^ step created artificial DNA sequences having extreme cyclizability that rarely exist in our dataset, likely because of deep network hallucination^39^ (Fig. 4c). During the selection, poly(dA:dT) tracts accumulated with lengths converging to 6 or 11 bp (half or single helical turn) that are positioned in phase along helical repeats (Supplementary Fig. 8) consistent with the model where phased repeats of (dA_5-6_:dT_5-6_) tracts can cause static bends to add up or cancel each other out depending on the relative phase^40–42^. We experimentally measured the kinetics of these sequences using single-molecule cyclization^4^ (Methods) and confirmed that the DNA sequence with the highest C0_corr_ rapidly looped (∼90% within 5 minutes), but the DNA with the lowest C0_corr_ hardly formed loops (∼10% after an hour, Fig. 4d). Atomic force microscopy (AFM) images of 600 bp sequences derived from *in silico* evolved sequences of the large positive and negative C0_corr_ values showed wavy structures and straight structures (Fig. 4e), respectively, with the corresponding curvature being much larger for the more cyclizable sequence (Fig. 4f).

To what extent do sequence-encoded DNA physical properties dictate protein-DNA interactions? It is important to reflect on the facts that cells have developed approaches to increase apparent DNA flexibility by the activities of both site-specific and site non-specific architectural DNA kinking proteins^43,44^. Proteins that contain one or more high-mobility group boxes (HMGB) are believed to play roles in DNA compaction. We therefore tested the DNA-bending ability of HMGB containing proteins, Nhp6A and HMGB1 (Methods). Remarkably, the extremely rigid DNA discovered via *in silico* evolution, which rarely cyclized even after 1 hour, showed greatly accelerated cyclization in the presence of Nhp6A or HMGB1 proteins (Fig. 4g). This result is an important reminder that architectural DNA-binding proteins can overcome intrinsic DNA physical properties.

### Eccentric mechanics of mitochondrial DNA of diplonemid protists

Despite the experimental validation of *in silico*-engineered DNA of extreme mechanics, we initially presumed that these artificial sequences, which required very strong selection over many rounds, would not appear in nature. Therefore, we were surprised when nucleotide BLAST^45^ search using the hallucinated sequence with the highest C0_corr_ found matches in the database of natural DNA sequences. Most of the matches were to the non-coding regions of mitochondrial genomes of *Namystynia karyoxenos* and *Hemistasia phaeocysticola*, both from Hemistasiidae, diplonemid protists living in the ocean (Supplementary Fig. 9a-c, Methods). The mitochondrial DNA (mtDNA) of diplonemids has a highly unusual architecture, with genes fragmented into small modules contained on different, non-catenated circular chromosomes that consist mostly of noncoding DNA. Transcription of these gene modules occurs independently, and after editing the transcripts are *trans*-spliced together, assembling the modules into mature mRNAs^46^. Sequence similarity between the non-coding region defines classes of mtDNA from A to Q, X, or U (unclassified)^46^.

We observed higher average cyclizability in the non-coding regions in the mtDNA of Hemistasiidae, compared to other diplonemids (Fig. 4h), which is due to the accumulation of poly(dA:dT) tracts in a specific pattern. For example, the highly cyclizable motif, 5’-GGGCCAAAAA- 3’, is present in the mtDNA of *H. phaeocysticola* with moderate frequency, while this motif greatly expanded its prevalence in *N. karyoxenos* (Supplementary Fig. 9d). This repeating motif was previously reported as an unorthodox characteristic that defines class X of mtDNA^46^. AFM imaging of 600 bp DNA derived from the mtDNA X031 of *N. karyoxenos* (Supplementary Fig. 9e, Supplementary Table 1) revealed wavy structures (Fig. 4e) with elevated curvature value (Fig. 4f), resembling the *in silico-* engineered sequence with extremely high cyclizability.

Unlike classical diplonemids that have reached a plateau in their complexity of gene fragmentation, hemistasiids have evolved additional gene fragmentation with twice the number of modules (Fig. 4h). Gene fragmentation, transcriptional, and post-transcriptional modification and regulation with substantial complexity may necessitate a large copy number of mtDNA as part of a mechanism to ensure the transmission of each module to subsequent generation without loss. Using transmission electron microscopy of *N. karyoxenos* we observed the presence of extraordinarily large amount of mtDNA, organized as strips of electron-dense beads within the organellar lumen (Fig. 4i). To quantify the total amount of mtDNA in this organism, we used propidium iodide (PI) because, unlike Hoechst 33342, its DNA straining is independent of DNA cyclizability (Supplementary Fig. 10a). Fluorescence intensity of PI staining after segmentation of nucleus volume from phase contrast images revealed that approximately 2/3 of total cell DNA is extra-nuclear (Fig. 4j, k, Methods). In the absence of nuclear genome size estimates for *N. karyoxenos*, we used the 280 Mb-haploid nuclear genome of the closely related *P. papillatum*^47^ as a proxy. This approximation led to an estimated ∼653 Mb mtDNA in the single mitochondrion of *N. karyoxenos*, which would qualify it as the largest amount of extra-nuclear DNA known so far with the previous record being approximately ∼260 Mb mtDNA^48^. To provide some context, in a typical human cell, numerous mitochondria combined contain ∼1000 times less DNA than the nucleus (∼8.3 Mb vs ∼6.4 Gb)^49^. We confirmed the abundant presence of highly cyclizable sequences in mitochondria by fluorescence *in situ* hybridization (Supplementary Fig. 10b).

In this study, we showed that proteins that contain HMGB make even the extremely rigid sequences cyclizable as rapid as the most cyclizable sequences (Fig. 4g), and HMGB proteins are found in the mitochondria of most organisms^50,51^, for example, Abf2p in yeast^52^ and TFAM in mammals^53^. Therefore, there may be no need for extremely cyclizable sequences as long as mtDNA compacting proteins exist. Publicly available genomic sequences of *N. karyoxenos* (https://www.ncbi.nlm.nih.gov/sra/SRX5472374 and https://www.ncbi.nlm.nih.gov/sra/SRX5434880) allow us to speculate which proteins implicated in mtDNA packaging are present. The search in the *de novo* assembly of the RNA-seq data^54^ did not identify any mtDNA-associated histone-like proteins (KAPs), which are the only known ones possibly involved in packaging mtDNA in these and related protists^55^. Extending our search to all other diplonemid sequences, including the high-quality nuclear genome of *P. papillatum*^47^, and using KAP1 through KAP4 of *Crithidia fasciculata* as queries failed to identify their putative homologs, allowing us to conclude that we could not identify any mtDNA binding proteins in diplonemids. Therefore, we suggest that the requirement to pack an extraordinarily large amount of DNA into a small volume of single mitochondrion in the absence of mtDNA compacting protein may have caused the accumulation of extremely cyclizable sequences in *N. karyxenos*. In fact, existing mutations around the 50 bp consensus sequence with extreme cyclizability rarely drop C0_corr_ below 2 in *N. karyoxenos* (only 0.15% of the time) whereas random mutations with the same Hamming distance from the consensus sequence can lower C0_corr_ below 1 (Supplementary Fig. 9f, Methods). Therefore, the native repeats have undergone mechanical selection to preserve high cyclizability of mtDNA.

To seek other examples of natural DNA of extreme mechanics, we ran BLAST search using a highly cyclizable mtDNA motif from *Artemidia motanka*, a close relative of *N. karyoxenos* and *H. phaeocysticola*^56^, and found matches to bird genomes, specifically their microchromosomes (see regions of house finch chromosome 39 and common parakeet chromosome 30 with cyclizability frequently exceeding 2) (Fig. 4l, Supplementary Fig. 11a). Of note, the closest outgroup, American alligator, did not show extreme cyclizability in its smallest chromosome (Supplementary Fig. 11b). Interestingly, we observed a strong anticorrelation between chromosome-averaged cyclizability and chromosome size in house finch (*Haemorhous mexicanus*) (Fig. 4m), and 59 other bird species (Fig. 4n) suggesting that there was a selection pressure to enrich for highly cyclizable sequences in tiny chromosomes in the bird lineage although we do not know the biological processes that the observed mechanical selection operated on. Overall, our discovery of extreme DNA mechanics in two unrelated lineages of uni-and multicellular eukaryotes of life suggests that our *in silico* method identified extreme biology and provides examples of nature finding a way to use even extremely eccentric DNA mechanics.

## Discussion

We significantly increased the accuracy of cyclizability prediction over prior methods by (1) removing DNA with low sequencing read counts, (2) mitigating the effect of adapter sequences, and (3) avoiding learning the cyclizability of DNA with Nt.BspQ1 nickase recognition motifs (Fig. 1, Methods). Predictions of DNA looping at a near-perfect accuracy will help link DNA mechanics to the function of genomic elements and understanding the evolution of DNA mechanics.

Phased (dA_5-6_:dT_5-6_) tracts accumulated during the *in silico* evolution (Supplementary Fig. 8) can explain the fluctuations of G/C contents or poly(dA:dT) tracts reported in genomic elements of many bacteria and eukaryotes where certain DNA mechanical properties are desired, e.g., *Saccharomyces* nucleosomes^10^, transcription start sites^57^, centromeres^34^, replication origins^58^, mtDNA of trypanosomes^59,60^, *E. coli* DNA gyrase cleavage sites^61^ and *Streptomyces* BldC binding motifs^12^.

Selection against mutations that disrupt the functionally important mechanical properties can in principle be quantified by examining mechanical features of sequences found in the genetic pool. Here, our investigation based on population genetics data of 1,011 yeast isolates identified their centromeres as a region with mechanical selection, likely due to unique mechanical properties under selection for inner kinetochore stability (Fig. 3). This introduces a new possibility to apply a similar method to diverse species, including humans, and relate DNA mechanics to phenotypes or diseases.

MtDNA occurs in nucleoids, and numerous nucleoid-associated proteins have so far been identified^50,51^. The consensus is that in the yeast mitochondrion, DNA is wrapped only by Abf2p^52^ thanks to its two HMGB boxes, each of which induces a sharp 90° bend^62^. The composition of mammalian mitochondrial nucleoid is still unclear, but TFAM, the HMGB boxes of which intercalate into the minor groove in sequence-unspecific manner^53^, compacts DNA by bending the DNA backbone and DNA loop formation until the DNA is fully compacted^63^. This is the same mechanism utilized by yeast Nhp6A and mammalian HMGB1 proteins tested here (Fig. 4g). The amplification of mtDNA in *N. karyoxenos* (Fig. 4i-l, Supplementary Fig. 10) and in other diplonemids^48^ may have required extreme mtDNA compaction, which can be achieved through DNA-bending proteins or by selection for DNA sequences with intrinsic properties that make them highly cyclizable. We propose that diplonemids evolved along the latter path; exploiting highly cyclizable mtDNA to store amplified mtDNA in the absence of DNA-bending proteins.

## Methods

### Uncertainty of loop-seq measured cyclizability

Variables were defined as follows. *N*, the total number of aligned reads of DNA molecules in a library, *n*, the number of aligned reads to a target DNA sequence, *C*, cyclizability. The lower-case c and s stand for control (no digestion) and sample (sequenced after digestion of unlooped DNA). Cyclizability determined by loop-seq is log((*n*_*s*_/*N*_*s*_)/(*n*_*c*_/*N*_*c*_)) (Supplementary Note 1).

Bayesian statistics with an uninformative Jeffreys prior yields a probability distribution of cyclizability.

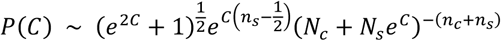

The 95% confidence interval (CI) of cyclizability is determined as *C_lower_* < *C* < *C_upper_*, where *P*(*C* < *C_lower_*) = 0.025 and *P*(*C_upper_* < *C*) = 0.025. Similarly, the uncertainty score calculated using frequentist statistics is as follows.

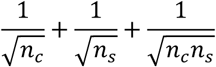

Here, we use, interchangeably, the 95% CI determined using Bayesian statistics and frequentist statistics (Spearman’s *R* = 0.997) to select a sub-dataset for model training and testing. See Supplementary Note 2 for details on the development of the formula.

### Model architecture

Models that share a single deep learning architecture (Supplementary Fig. 3a) were trained individually on different cyclizabilities (C0, C26, C29, and C31 etc. where the numeric values denote the positions of biotin used for tethering DNA molecules to a bead surface). The models were implemented using Keras^64^.

Input (200,) - A 50 bp DNA is converted into a 200-dimensional vector by one-hot encoding:

A: [1, 0, 0, 0], T: [0, 1, 0, 0], G: [0, 0, 1, 0], C: [0, 0, 0, 1]

First 1D convolution layer - Kernel size: 28, Output channels: 64, Stride: 4, Output shape: (44, 64). A bias term and a rectified linear unit (ReLU) activation were added.

Second 1D convolution layer - Kernel size: 33, Output channels: 32, Output shape: (12, 32). A bias term and a rectified linear unit (ReLU) activation were added.

Flatten layer - Output shape: (384,).

First fully connected layer - Output shape: (50,). A bias term and a rectified linear unit (ReLU) activation were added. The layer was L2 regularized with the regularization constant of 0.001.

Second fully connected layer (output) - Output shape: (1,). This layer predicts cyclizability of a 50 bp DNA sequence.

### Training the model

Training datasets for C26, C29, or C31 consist of sequences from the Tiling library^5^ selected by their uncertainty score of measured C values lower than 0.1. C0 was trained using sequences that are shared in the three datasets used for training C26, C29, and C31. The testing datasets were selected in the same way from sequences in the ChrV library^5^. Sequences containing the digestion motifs of the endonuclease Nt.BspQ1, 5’-GAAGAGC-3’ or 5’- GCTCTTC-3’, were removed from all datasets, because the unintended nicks produced in the variable 50 bp DNA during the loop-seq protocol increase C values. The effect of the digestion motif on cyclizability was previously reported and interpreted erroneously as being caused by changes in DNA mechanics^7^. Unless otherwise stated, models were trained to minimize a mean squared error loss for seven epochs using the Adam optimizer^65^, with an initial learning rate of 0.001 and decay rates β_1_ and β_2_ of 0.9 and 0.999, respectively. The computational details are described in Supplementary Note 3.

### Removing the effects of adapter sequences

The upper envelopes *U*(*n*) of the oscillatory patterns of *C*(*n*) on a sufficiently long DNA were acquired by cubic interpolations of local maxima using the interp1d function of Python SciPy 1.9.3^66^ (Supplementary Fig. 5), where *n* is the position of biotin tether (26, 29, or 31). A cyclizability value is considered a local maximum if the value is the highest among a set of 7 consecutive cyclizability values (including itself, three values to the left and three values to the right). The process is similarly done to find lower envelopes *L*(*n*).

The corrected upper envelope *U*0 and the mock amplitude *A*′ (in which the difference between *U*(*n*) is absorbed) that fits to the formula below are calculated using the fsolve function of Python SciPy 1.9.3^66^ (Supplementary Fig. 5a).

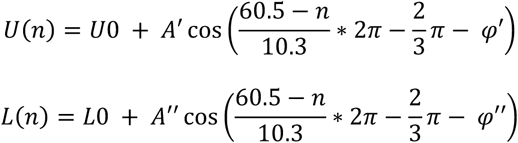

The adapter-corrected cyclizability is defined accordingly.

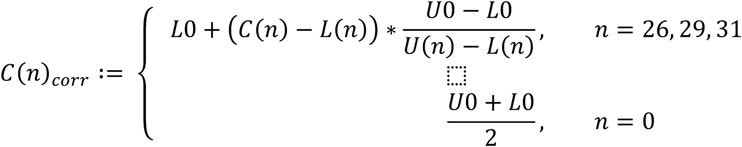

C0_corr_, C26_corr_, C29_corr_, and C31_corr_ were calculated at a base resolution for yeast ChrV. We selected 576,647 50 bp windows with clearly defined adapter-corrected cyclizability values. This selection excluded the first and last 50 windows, as well as any windows with a corrected upper envelope lower than the corrected lower envelope. The selected data were used for learning over four epochs with the same model hyperparameters stated in the previous section (Supplementary Fig. 3a). The model predicted adapter-corrected cyclizability with high accuracies in testing datasets that are not used for model training (Pearson’s R > 0.96, Supplementary Fig. 5e). The computational details are described in Supplementary Note 4.

### Aligning genomic sequences of 1,011 yeast isolates

We aligned the genomic sequences of 1,011 isolates of *S. cerevisiae* ^25^ to a region of interest using the BLAST-like alignment tool (BLAT)^67^. Template DNA sequences were obtained from the sacCer3 reference genome^68^. Any alignments with less than 80% identity to the reference, as well as those containing ambiguous nucleotides or indel mutations, were excluded. To avoid including excessively duplicated regions in our dataset, we excluded genomic regions with more than 1.2×1,011 alignments, in accordance with the process outlined in a previous study^24^. Aligning the genomic sequences in centromeres is outlined in Supplementary Note 7.

### Quantifying selection pressure on DNA mechanics

We quantified mechanical selection based on population genomes aligned to a region of interest^25^. For each 50 bp window, simulated sequences were generated from the natural alignments with random mutations, such that Hamming distance from the consensus sequence is identical in both sets (Fig. 3a). Hamming distance of a 50 bp DNA is the minimal number of point mutations required to create that 50 bp DNA from the consensus sequence, and the consensus sequence is the DNA sequence in which the most frequent base at each position is taken as the consensus. Natural or simulated sequences identical to the consensus sequence were omitted from the analysis. An identical mutant generating scheme was used to study the effect of native mutations in *N. karyoxenos* mtDNA (Supplementary Fig. 9f).

We predicted C0_corr_ of natural and simulated 50 bp sequences. As yeast isolates share a common ancestor, natural alignments consist of groups of homogeneous 50 bp sequences. Accordingly, we set the variables as follows, assuming that there are *n* different natural C0_corr_ values (*i* is an integer ranging from 1 to *n*). *c*_*i*_: Counts of natural isolates with identical DNA sequence *i*. *N*: Total counts of simulated sequences. *r*_*i*_: Rank of each natural C0_corr_ among simulated 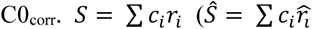 for observed ranks). The rank of 0 is assigned to a natural sequence if it has the lowest C0_corr_ among simulated C0_corr_.

As an analogue of *Z* value from normal distribution, *Z*-score of mechanical selection is determined.

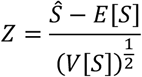

where 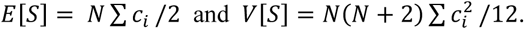

*P*-value of mechanical selection is determined as follows.

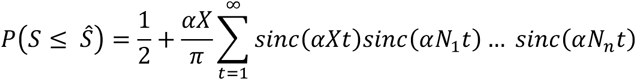

where 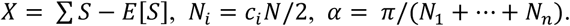

Here, *sinc* is defined as follows.

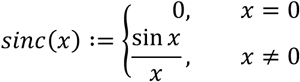

The statistics are valid only in 50 bp windows with enough diversity of natural sequences (Supplementary Note 6). We measured the diversity using information entropy.

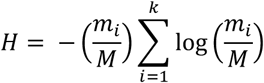

*M* is the total number of natural sequences, with *k* unique sequences each with *m*_*i*_(*i* = 1,…, *k*) isolates (*M = ∑ m*_*i*_). 50 bp windows with information entropy higher than 0.75 were used in further analyses. Reproducibility and computational details are described in Supplementary Note 6.

### Spatial analysis of DNA bending

C26_corr_, C29_corr_, and C31_corr_ measure proficiencies of DNA looping in three different directions^5^. Taking advantage of this, we inferred the rotational phase of DNA looping and its amplitude using a method that we named spatial analysis.

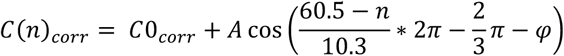

*n* is the position of biotin tether (26, 29, or 31). The values of C26_corr_, C29_corr_, and C31_corr_, were adjusted by subtracting constants to set the average of each cyclizability to 0 before use, because cyclizability values from different loop-seq experiments can be compared up to a constant offset^5^. C0_corr_, amplitude *A* ≥ 0, and phase φ ∈ [−π, π), were then obtained using Python SciPy 1.9.3 package^66^ with the starting estimate of [1, 1, 1]. The amplitude *A* is a relative preference of bending in a certain direction indicated by the phase φ. For example, a DNA molecule has no preference in bending direction when the amplitude is zero.

The formula assumes 10.3 bp for a helical turn of a duplex DNA, but the precise value may vary in different contexts. Thus, when interpreting the results, we avoid relying excessively on the precise values of amplitude *A* or phase φ. For a similar reason, C0_corr_ obtained by spatial analysis was not used in model training or testing. Verification of the method is described in Supplementary Note 5.

We repeat the process using *Z*-scores to see the mechanical selection in a 3-dimensional space.

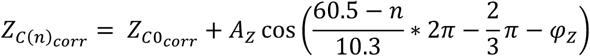

*Z*_*C*(*n*)_*_corr_* are *Z*-score obtained by using C(n)_corr_ instead of C0_corr_, where *n* is the position of biotin tether (26, 29, or 31). *A*_*Z*_ and φ_*Z*_quantify the amplitude and the rotational phase of mechanical selection in a 3-dimensional space, respectively. Computations to obtain *A*_*Z*_and φ_*Z*_ using the formula above are done similarly as in the spatial analysis based on cyclizability. Unlike cyclizability, the *Z*-scores were not normalized by subtracting their average values before analysis, as the constant offsets between different types of cyclizability disappear during *Z*- score computation.

### Visualizing DNA bending in a 3-dimensional space

The main helical axis of a dsDNA molecule is defined by the midpoint of C6 of pyrimidine and C8 of purine base^69^. To visualize the intrinsic DNA bending at a base-base step, we drew an arrow perpendicular to the main helical axis. The starting point of each arrow was set to the midpoint of the helical axis at a base-base step, and the length (in angstroms) of each arrow represented the amplitude of DNA bending (obtained by spatial analysis) multiplied by a factor of 30 unless otherwise noted. For each base-base step, a spatial analysis result from the 50 bp window that puts the base-base step in the middle (25 bp to the upstream and downstream) was used. For a base-base step at the boundaries of a linear DNA, the average cyclizability of 200 DNA sequences with randomly filled missing bases was used for spatial analysis.

The observed bending vector of a base-base step was defined by the difference between the unit vector representing the direction of main helical axis at the last (between the 49^th^ and 50^th^ bases) and at the first (between the 1^st^ and 2^nd^ bases) base-base step of the surrounding 50 bp window. The similarity between the intrinsic and observed DNA bending was defined by the inner product of the vectors representing the intrinsic and the observed bending. PDB structures generated by refining and fitting experimental data to other known PDB structures were not included in our analysis at the RCSB database scale (Fig. 2) to avoid arbitrary DNA sequences being inserted into the structures.

### Total internal fluorescent microscopy slide preparation for single molecule colocalization analysis of centromere assembly

Coverslips and microscope slides were ultrasonically cleaned and passivated with PEG as described previously^35^. Briefly, slides were ultrasonically cleaned and then treated with vectabond (Vector Laboratories) prior to incubation with resuspended 1% (w/v%) biotinylated mPEG-SVA MW-5000K/mPEG-SVA MW-5000K (Lysan Bio) in flow chambers made with double-sided tape. Passivation/functionalization was carried out overnight at 4 °C. After functionalization, flow chambers were washed with Buffer L (25 mM HEPES pH 7.6, 2 mM MgCl_2_, 0.1 mM EDTA pH 7.6, 0.5 mM EGTA pH 7.6, 0.1 % NP-40, 175 mM K-Glutamate, and 15% Glycerol) and then incubated with 0.3 M BSA/0.3M Kappa Casein in Buffer L for 5 min. Flow chambers were washed with Buffer L and then incubated with 0.3M Avidin DN (Vector Laboratories) for 5 min. Flow chambers were then washed with Buffer L and incubated with ∼100 pM of respective CEN DNA template (Supplementary Table 1) for 5 min and washed with Buffer L. For endpoint colocalization assays, flow chambers were filled with 100 μL of whole-cell extract (WCE) containing protein(s) of interest via pipetting and wicking with filter paper. WCE was prepared as previously described^35^. After addition of WCE, slides were incubated for 90 min at 25°C and then WCE was washed away with Buffer L. Flow chambers were then filled with Buffer L with oxygen scavenger system^70^ (10 nM PCD/2.5 mM PCA/1mM Trolox) for imaging.

### Total internal fluorescent image collection and analysis

Colocalization images were collected on a Nikon TE-2000 inverted RING-TIRF microscope with a 100× oil immersion objective (Nikon Instruments) with an Andor iXon X3 DU-897 EMCCD camera. Images were acquired at 512 px × 512 px with a pixel size of 0.11 µm/px at 10MHz. Atto-647 labeled CEN DNA templates (Supplementary Table 1) were excited at 640 nm for 300 ms, GFP-tagged Cse4 was excited at 488 nm for 200 ms, and mCherry-tagged Ndc10 was excited at 561 nm for 200 ms. Single snapshots of all channels were acquired, and images were analyzed using ComDet v.0.5.5 plugin for ImageJ (https://github.com/UU-cellbiology/ComDet) to determine colocalization and quantification between DNA channel (647 nm) and GFP (488 nm) and mCherry (561 nm) channels. Results were quantified and plotted using MATLAB (The Mathworks, Natick, MA). Adjustments to example images (contrast, false color, etc.) were made using FIJI^71^ and applied equally across entire field of view of each image.

### Local alignment searching of DNA sequences with high cyclizability

Matches of 5’-GCCAAAAAAGGGCCAAAAATGGCCATTTTTGGCCCTTTTTTGGCCTTTTT-3’, the 50 bp DNA with the highest C0_corr_ found after the 50^th^ steps of *in silico* selections favoring higher C0_corr_ (Fig. 4c, Supplementary Fig. 8e), were found using BLASTn search with the word size, match, and mismatch scores of 7, 1, and-1, respectively. Resulting hits were sorted by E-value (Supplementary Fig. 9a, b). C0_corr_ of mitochondrial genomes, including those found by BLASTn search, were predicted after replacing ambiguous nucleotides with random bases (Fig. 4h, Supplementary Fig. 9c).

### Ultrastructure of Namystynia karyoxenos

*N. karyoxenos* was cultivated in a nutrient-rich medium at 22 °C as previously described^56^. The cells were harvested during the exponential growth phase by centrifugation at 4,000 g for 30 min and then processed by high-pressure freezing technique and freeze substitution as described elsewhere^72^. Subsequently, the samples were observed using a JEOL 1400 transmission electron microscope at an accelerating voltage of 80 kV.

### Quantification of mitochondrial DNA

Cells were harvested as described above and fixed with 4% paraformaldehyde in artificial seawater for 30 min at room temperature (RT). After fixation, the paraformaldehyde was washed off with phosphate buffered saline (PBS), and the cells were mounted onto gelatin-coated slides for adhesion. The air-dried slides were then immersed in −20 °C methanol overnight for cell permeabilization, following which the cells were rehydrated in PBS for 10 min and treated with RNAse A (50 μg/mL) for 2 hrs at RT. Cells were then stained with a combination of 5 µg/ml Hoechst 33342 (bisbenzimide) and 25 µg/ml propidium iodide (PI) for 10 min. Dyes were removed by a wash in PBS and slides were mounted with ProLong Gold Antifade reagent (Invitrogen). Images were acquired via a 100× objective lens on a BX63 Olympus widefield fluorescence microscope equipped with an Olympus DP74 digital camera using CellSens Dimension software v. 1.11 (Olympus) and processed in Image J v. 1.52p software. From micrographs of the PI fluorescence we first measured the background signal from the modal pixel value of the field of view and subtracted it from the pixel values. We then identified and segmented the nucleus, based on their constant size and internal structure in the PI image, and whole cell, from the phase contrast image. The relative nuclear to total DNA quantity was measured from the ratio of integrated pixel values in the nucleus and whole cell regions. As the cells are thin, we used widefield microscopy and a single focal plane, which effectively integrates the fluorescent signal from the entire cell volume.

For in vitro measurement of Hoechst 33342 and PI fluorescent signal when bound to DNA of different cyclizabilities, we generated double stranded (ds) DNA by annealing of a forward and reverse 100 base oligonucleotides. 100 nM forward and reverse primer were mixed in annealing buffer (10 mM Tris-HCl, pH 8.0; 50 mM NaCl; 1 mM EDTA), and annealed by denaturation at 95°C for 2 min, followed by cooling to 25°C over 45 min. The dsDNA was diluted to 25 mM in annealing buffer with 5 μg/ml Hoechst 33342 and 25 μg/ml PI, incubated at RT for 15 min, and fluorescence was measured at 544 nm excitation/620 nm emission (red fluorescence) and 355 nm excitation/460 nm emission (blue fluorescence). Background signal from DNA-lacking sample was measured and subtracted. All samples were generated and measured in technical triplicate.

### Targeting the circularized regions of *N. karyoxenos* mtDNA

To detect circularized mtDNA sequence, we employed fluorescence in situ hybridization (FISH) in combination with immunofluorescence (IF) assay to visualize the mitochondrion. The cells were fixed as described above, washed and permeabilized using eBioscience buffer, and incubated overnight at 4° C with a rabbit antibody against β chain of mitochondrial ATP synthase, diluted 1:500. The primary antibody was then removed, and the cells were washed and incubated with a goat anti-rabbit secondary antibody conjugated to Alexa Fluor 488, diluted 1:1,000, for 1 hour at RT in the dark. Next, the cells were washed and allowed to adhere onto gelatin-coated slides, while being kept in the dark. The air-dried slides were then treated with 0.1% Triton X-100 for 5 min and washed with PBS. Following this, the cells were infiltrated with a DNA probe (5’-Cy3-CCAAAAAAGGGCCAAAAATGGCC-3’) that was resuspended in a hybridization buffer (70% formamide; 1× saline sodium citrate (SSC) buffer pH 7.0; 10% dextran sulfate; 8 µg salmon sperm DNA; 50 ng DNA-Cy3 probe) for 1 hr at RT. The samples were then denatured for 5 min at 85 °C and left for hybridization overnight in the dark at 42 °C in a humid chamber. Afterwards, the samples were washed twice for 15 min with 70% formamide and 10 mM Tris-HCl, pH 7.2, at 42 °C and then 3× for 5 min with 1× SSC. Subsequently, the slides were mounted with ProLong Gold antifade reagent (Invitrogen) containing Hoechst 33342 and examined by FV3000 confocal laser scanning microscope (Olympus) with the spectral filter windows set as follows: for Hoechst channel 417-486 nm, Alexa Fluor 488 505-537 nm, and for Cy3 549-584 nm.

### Single-molecule fluorescence resonance energy transfer (smFRET) DNA cyclization assay

The instrumental setup of the single-molecule total internal reflection fluorescence (smTIRF) microscope has been previously described^73^. The DNA constructs designed for the smFRET DNA cyclization assay are detailed in Supplementary Table 1. Single-stranded DNA labeled with fluorophores and biotin was purchased from Integrated DNA Technologies (IDT). For annealing, complementary single-stranded DNAs were resuspended in nuclease-free duplex buffer (IDT), mixed at a 1:1 molar ratio, heated to 95°C for 2 minutes, and then cooled to 25°C for over an hour. Polyethylene glycol (PEG)-passivated quartz slides were prepared and assembled according to established protocols^74^. For smTIRF imaging, biotin-labeled DNA oligos were diluted to 50 pM in T10 buffer (10 mM Tris-HCl (pH 8.0), 10 mM NaCl) and heated at 55 °C for 5 minutes to prevent premature annealing via sticky ends before immobilizing on the quartz slide. The DNA was immobilized on the quartz slides through biotin-neutravidin interaction and subsequently washed with T10 buffer after a 2-minute incubation.

In the real-time cyclization assay (Fig. 4d), a salt-free imaging buffer (100 mM Tris-HCl (pH 8.0), saturated Trolox, 8 mg/ml Dextrose, 0.83 mg/ml glucose oxidase, and 20 U/ml catalase) was introduced to keep the DNA oligos unlooped prior to imaging. For the real-time cyclization experiments, a high-salt imaging buffer (100 mM Tris-HCl (pH 8.0), 1 M NaCl, saturated Trolox, 8 mg/ml Dextrose, 0.83 mg/ml glucose oxidase, and 20 U/ml catalase) was flowed into the channel by a syringe pump. At designated time points, a 20-frame short movie was recorded (10 frames of green excitation, 10 frames of red excitation, 100 ms exposure time). If the imaging time point exceeded 30 minutes after the imaging buffer was flowed into the channel, fresh imaging buffer with the identical salt concentration was introduced into the channel to avoid low pH conditions caused by the oxygen scavenging system. The movies were analyzed using smCamera software^73,75^, selecting molecules with both green and red emissions for plotting the FRET efficiency histograms. For each time point, ∼1000 molecules were used to plot the FRET histogram. The fraction of unlooped and looped DNA oligos were quantified by fitting Gaussian distributions to the low and high FRET peaks using OriginLab software.

For measuring protein-induced DNA cyclization (Fig. 4g), the low C0_corr_ DNA (Supplementary Table 1) was immobilized on the quartz slide using the same protocol described above. The protein binding imaging buffer used was 20 mM Tris (pH 7.5), 150 mM NaCl, 1.5 mM MgCl2, 0.5 mg/ml BSA, saturated Trolox, 8 mg/ml Dextrose, 0.83 mg/ml glucose oxidase, and 20 U/ml catalase. The low C0_corr_ DNA was equilibrated in the protein binding buffer in the channel for 30 minutes, and short movies were recorded to confirm the initial FRET distribution in the absence of proteins. The desired protein was diluted to approximately 60 nM in the protein binding imaging buffer and flowed into the channel using a syringe pump. A series of 20-frame short movies were recorded at designated time points. Data analysis and looping kinetics were conducted in the same manner as described above. All single-molecule measurements were performed at room temperature (∼22°C).

### AFM Sample Preparation

All DNA oligonucleotides with defined cyclizability were purchased from Ansa Biotechnologies, a DNA manufacturer with expertise in producing long repetitive sequences (Supplementary Table 1). A freshly cleaved mica surface was coated with 20 mM MgCl₂ buffer for 5 minutes, followed by two washes with distilled water. DNA samples, incubated in Tris-HCl buffer with 20 mM MgCl₂ and diluted to 0.1 nM, were then deposited onto the freshly cleaved, MgCl₂-coated mica surface. After a 5-minute incubation, samples were rinsed with either distilled water or MgCl₂ buffer and subsequently imaged using high-speed AFM.

### Specification of High-Speed Atomic Force Microscope (HS-AFM)

Experiments on DNA cyclizability across different sequences were conducted using a commercial Sample-Scanning High-Speed Atomic Force Microscope (SS-NEX Ando model) from RIBM (Research Institute of Biomolecule Metrology Co., Ltd.). The AFM was operated in tapping mode to minimize interference with the deposited sample, and all samples shown in this paper were imaged in solution. Ultra-Short Cantilevers (USC- F1.2-k0.15-10), specifically designed for high-speed AFM, were employed with a resonance frequency of 1200 kHz, a spring constant of 0.15 N/m, and a length of 7 µm. These cantilevers were purchased from NanoAndMore. A wide scanner was used, with scan speeds ranging from 0.05 to 1 frame per second and resolutions set between 200 × 200 and 500 × 500 pixels.

### Image Processing and Analysis

HS-AFM images were viewed and analyzed using Kodec 4.4.7.39 software (Sakashita M, M Imai, N Kodera, D Maruyama, H Watanabe, Y Moriguchi, and T Ando. 2013. Kodec4.4.7.39). All images were processed with an X- Resonance Noise Filter and X lineTilt correction; detailed image correction protocols are described in the literature^76^. Contrast adjustments were applied to enhance structural features in the images. The length and the average curvature of DNA molecules were analyzed using Fiji^71^. DNA molecules shorter than 160 nm were excluded from the analysis due to the possibility of incomplete synthesis or digestion.

## Data availability

All adapter-dependent cyclizability measurements were downloaded from the sequencing data deposited in NCBI Sequence Read Archive under accession number PRJNA667271, and the genomes of 1,011 yeast isolates were obtained from accession number ERP014555. The mitochondrial genomes of *N. karyoxenos*, *H. phaeocysticola*, *A. motanka*, *S. specki*, *L. lanifica*, *R. humris*, *D. ambulator*, and *D. japonicum* were obtained from the NCBI Nucleotide database under accession number MN109419-MN109581, LC114082-LC114083, MN109174- MN109319, MN109336-MN109400, MN108931-MN109016, MN109083-MN109155, MF436742-MF436795, and MN109036-MN108966. Nucleosome occupancy data through chemical cleavage around nucleosome dyads in *S. cerevisiae* was obtained under NCBI Gene Expression Omnibus accession number GSE97290.

## Code availability

Code is available on GitHub at https://github.com/codergirl1106/Cyclizability-Prediction-Website. A web app is available at https://cyclizability-prediction-website-5vbkhabttypl6n29hkxc8q.streamlit.app/.

## Acknowledgements

J.P. and T.H. thank Martha L. Bulyk, Luca Marinai, and Xiao Liu for their advice on the applications of the model. J.P. and T.H. thank Yaojun Zhang, Margarita Gordiychuk, Bohao Fang, and Scott V. Edwards for their helpful discussion. J.P. thanks Basilio C. Huaman for his advice on deep-learning models. This work was supported by the National Institutes of Health grants (R35 GM143949 to L.J.M., R35 GM134842 to C.L.A, R35 GM149357 to S.B., R35 GM122569 to T.H.), the Czech Grant Agency (23-06479X to J.L.) and Wellcome Trust Sir Henry Dale Fellowship (211075/Z/18/Z to R.J.W.). S.B. and T.H. are Investigators of the Howard Hughes Medical Institute.

## Author contributions

J.P. and T.H. designed the research. J.P. performed all aspects of the research and data analysis. J.P. and T.H. wrote the paper. Other authors contributed to the following areas: G.P. obtained microscopy images of *N. karyoxenos*. A.R.P., D.J.H., and J.D.L. conducted single-molecule TIRF experiments and measured the assembly of inner kinetochore. J.H. conducted single-molecule FRET DNA cyclization assay. T-W.L. conducted atomic force microscopy. S.Y. developed the web prediction tool and verified analysis results for protein-DNA complexes collected from RCSB database. B.K.W. developed heuristic methods to accelerate cyclizability predictions. N.A.B. and L.J.M. provided Nhp6A and HMGB1 for DNA cyclization assay. A.B. advised on the analysis of loop-seq datasets. R.J.W. quantified the content of mtDNA in *N. karyoxenos*. A.B., L.J.M., R.J.W., C.L.A., S.B., and J.L. provided helpful scientific discussion and supported scientific collaboration. All authors commented on the manuscript.

---

**Supplementary Fig. 1.**
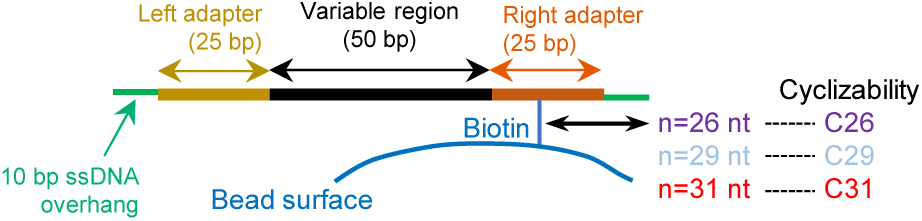
Schematic of a typical DNA molecule in loop-seq. DNA molecule for loop-seq consists of a variable region (50 bp) surrounded by two adapters (25 bp each) and 5’ ssDNA overhangs (10 nt). The ssDNA overhangs are complementary to each other and form dsDNA after looping^5^. DNA looping (or cyclization) reaction rate depends on biotin position for sequences that induce static bending, and the biotin position dependence of cyclizability can be eliminated by performing loop-seq with three different biotin positions indicated yielding C26, C29 and C31, and mathematically correcting for the position effect (Methods).

**Supplementary Fig. 2.**
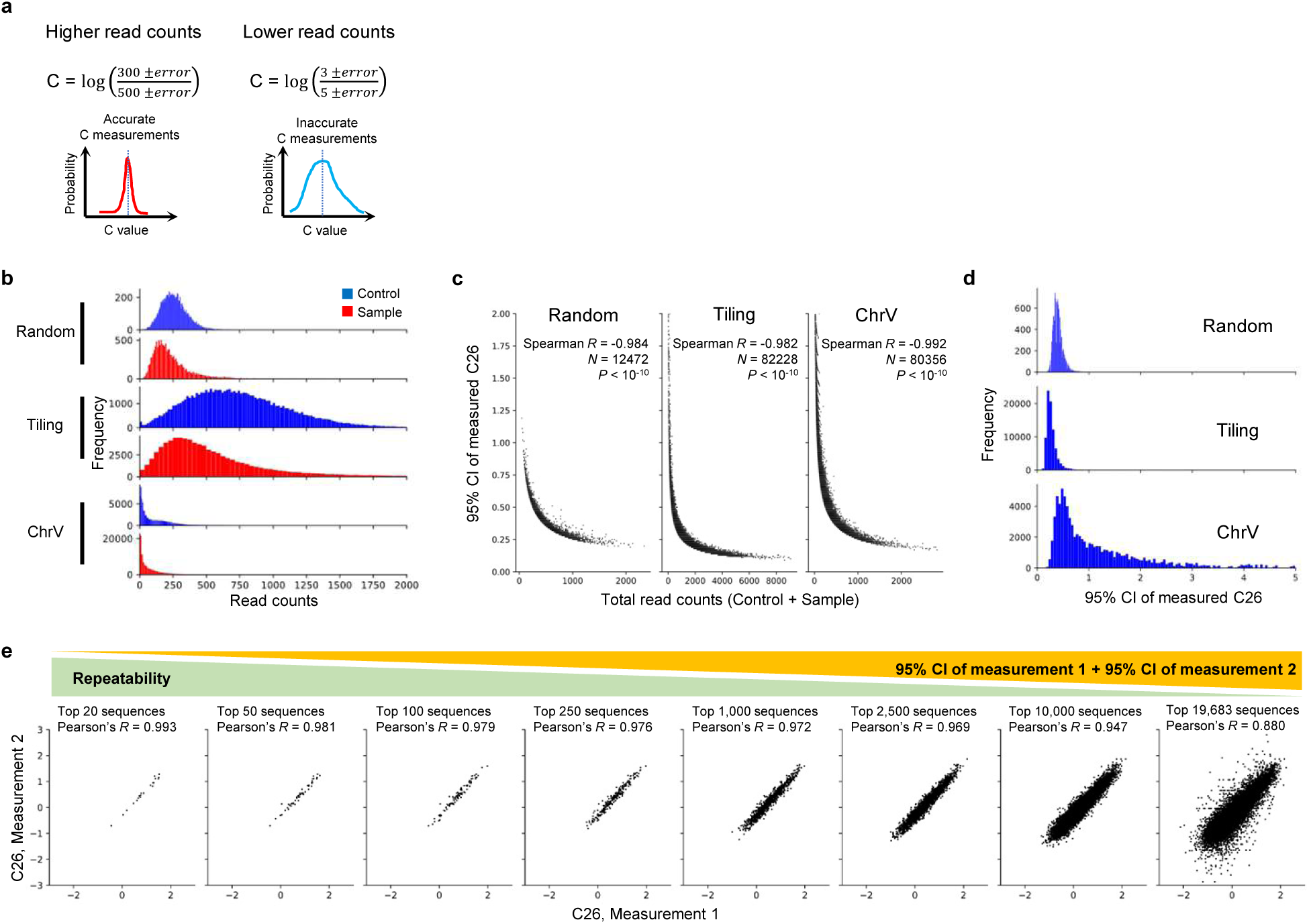
Read counts are anti-correlated with the 95% CI of measured cyclizability. **a**, Read counts vs measured cyclizability distribution. Lower read counts would give rise to a broader distribution because relative errors are larger. **b**, Distribution of read counts in loop-seq experiments measuring C26 in the Random, Tiling, and ChrV library. **c**, Total read count (before + after digestion) of each sequence vs the corresponding 95% CI of measured C26 in Random, Tiling, and ChrV library. **d**, Distribution of the 95% CI of C26 in the Random, Tiling, and ChrV library. **e**, Scatter plots of repeated C26 measurements on the Cerevisiae Nucleosome library. Top 20, 50, 100, 250, 1,000, 2,500, 10,000, 19,638 sequences with the lowest sum of 95% CI of repeated measurements were selected. Sample sizes and Pearson’s correlations are shown. Measurement 1 is from loop-seq of the mixture of Random and Cerevisiae Nucleosome library. Measurement 2 is from the 1- minute time point of the timecourse loop-seq on the Cerevisiae Nucleosome library.

**Supplementary Fig. 3.**
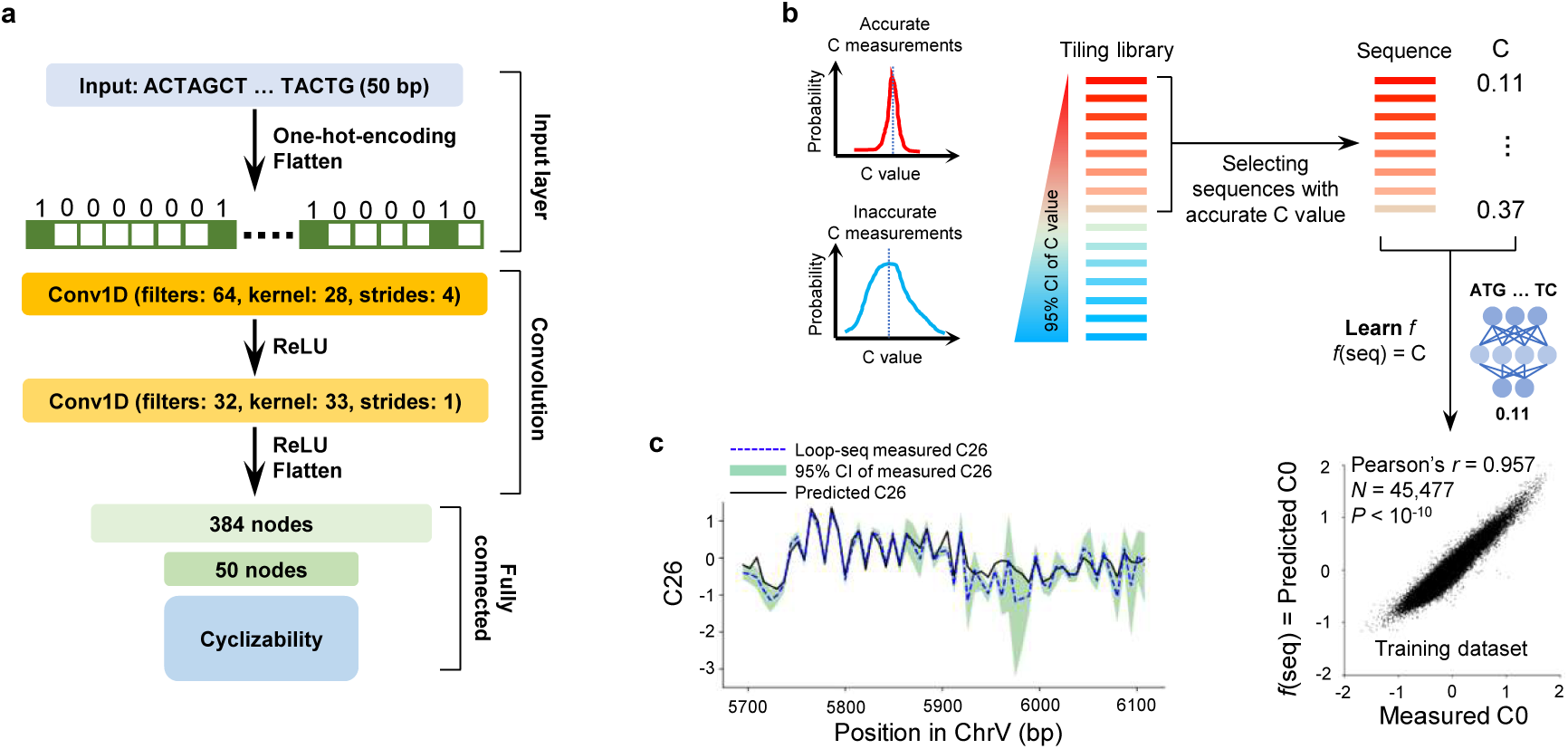
Training model to predict cyclizability. **a**, Model structure for learning cyclizability. **b**, Schematic of the model training process and scatter plot showing measured vs predicted C0 for the training dataset curated from the Tiling library^5^. Pearson’s correlation, sample size, and the corresponding two-tailed *P*-value are shown. Note that we are showing the same scatter plot as in Fig. 1b in this expanded schematic of model training. **c**, Measured C26, 95% CI, and predictions for an example region of yeast chromosome V.

**Supplementary Fig. 4.**
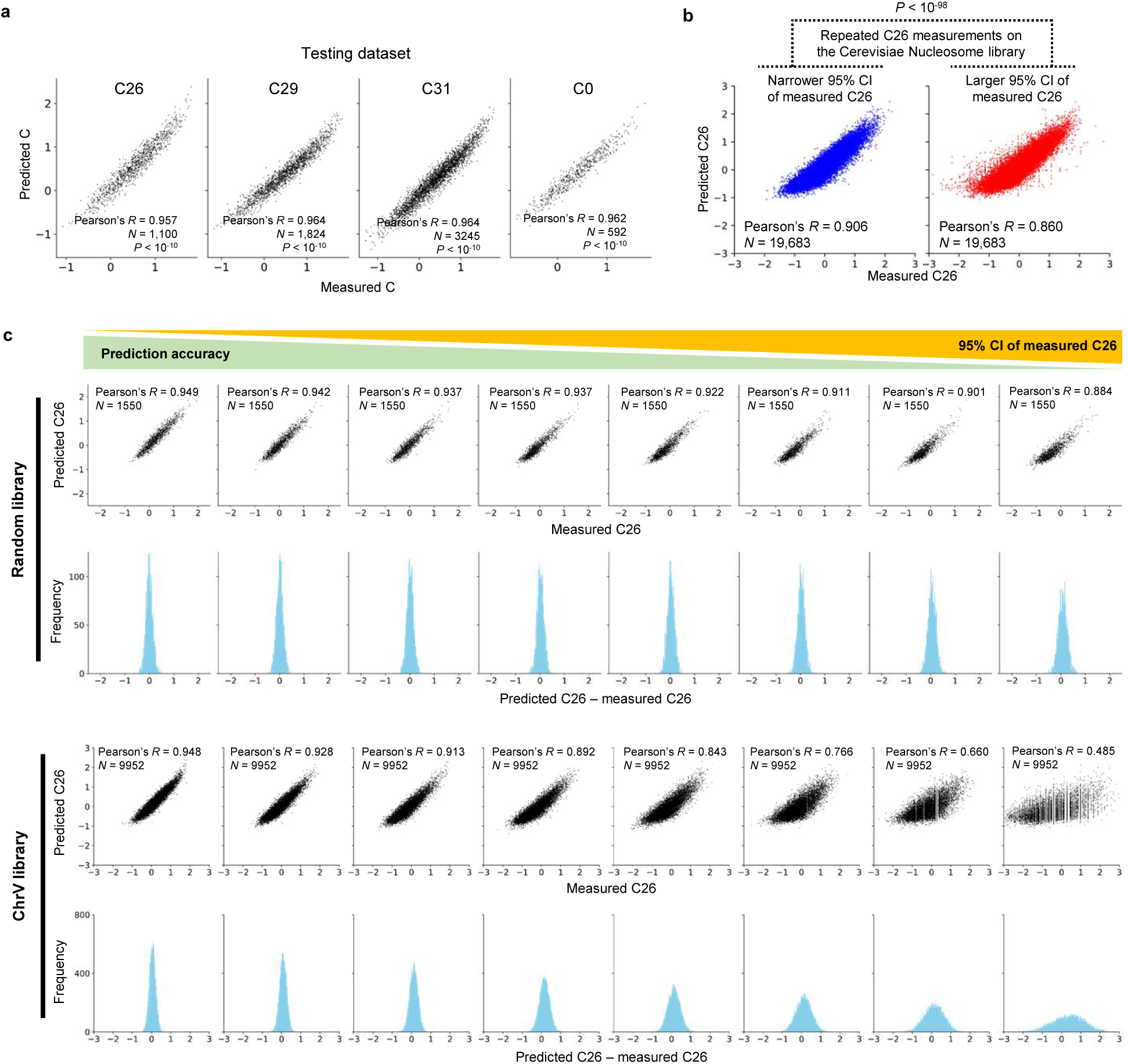
Prediction accuracy of cyclizability is affected by read counts. **a**, Measured vs predicted cyclizability of sequences with an uncertainty score below 0.1 in the ChrV library. **b**, Repeated measurements of C26 for the Cerevisiae Nucleosome library compared to predictions. For each sequence, the C26 measurement with the narrower 95% CI is used in the left plot (Pearson’s *R* = 0.906), while the measurement with the wider 95% CI is used in the right plot (Pearson’s *R* = 0.860). C26 measurements with narrower 95% CI show a stronger correlation with the predicted C26 (*P* < 10^-98^). For the repeated measurements, the mixture of Random and Cerevisiae Nucleosome library and timecourse loop-seq on Cerevisiae Nucleosome library at 1-minute were compared. **c**, Measured vs predicted C26 in subgroups of the Random and ChrV library. Sequences were sorted and classified into 8 subgroups based on the width of 95% CI of C26.

**Supplementary Fig. 5.**
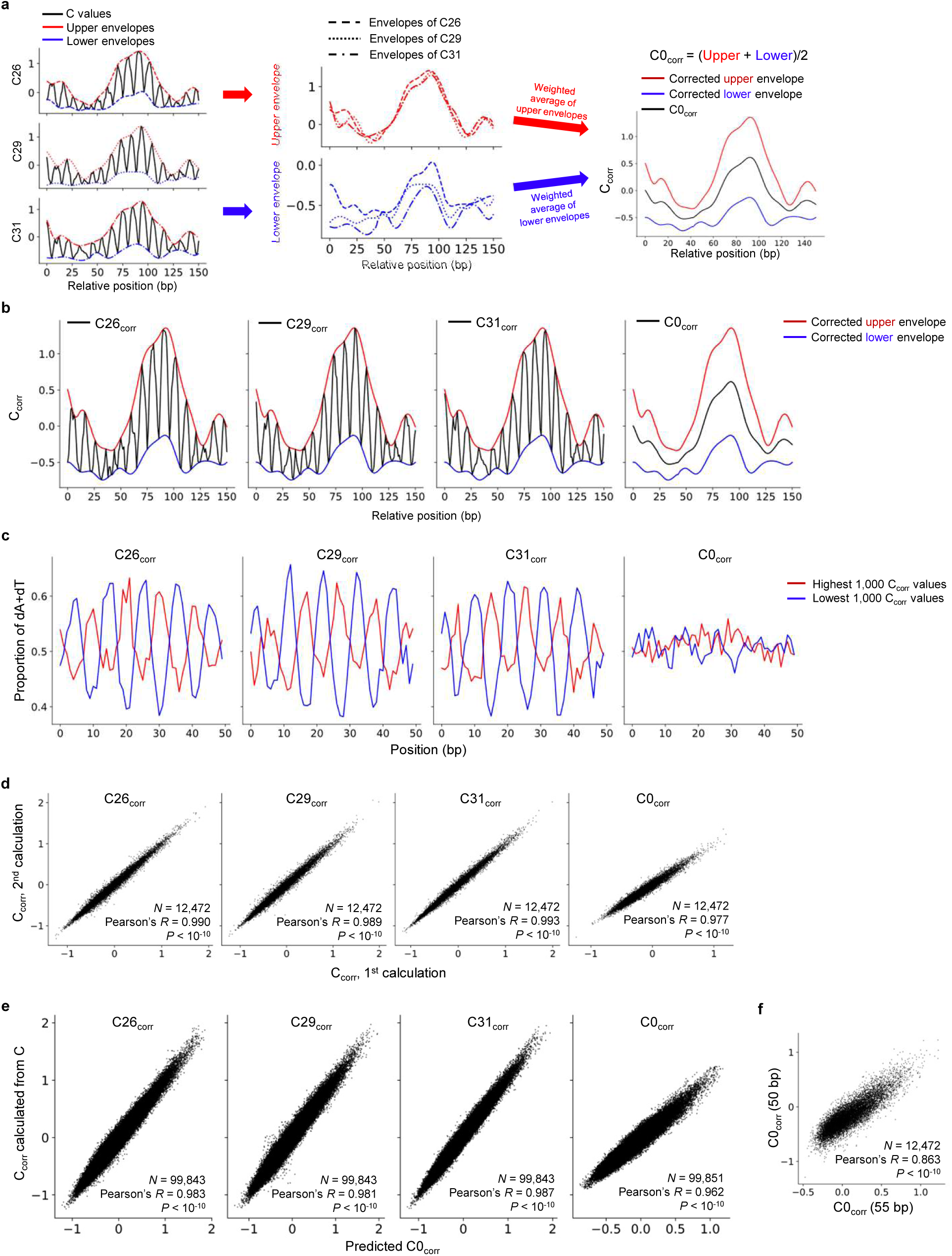
Defining and learning adapter-corrected cyclizability. **a**, Schematic of the procedure for defining the corrected upper and lower envelopes, and C0_corr_ (Method). **b**, The relationship between adapter-corrected cyclizability and corrected envelopes. **c**, AT content across 50 bp DNA averaged over 1,000 sequences curated from the Random library with the highest C_corr_ values, red, or the lowest C_corr_ values, blue. **d**, Scatter plots comparing repeated calculations of C_corr_. Different sets of upstream and downstream 50 bp sequences were used for repeated calculations of C_corr_ (Supplementary Note 4) **e**, C_corr_ calculated from predicted C values vs the corresponding C_corr_ values predicted directly from models that are trained using C_corr_. **f**, C0_corr_ of 55 bp DNA sequences in the library L vs the corresponding C0_corr_ of 50 bp DNA sequences in the Random library. A set of sequences in the library L and the Random library share the same 50 bp from the 5’ end. Details can be found in Supplementary Note 4.

**Supplementary Fig. 6.**
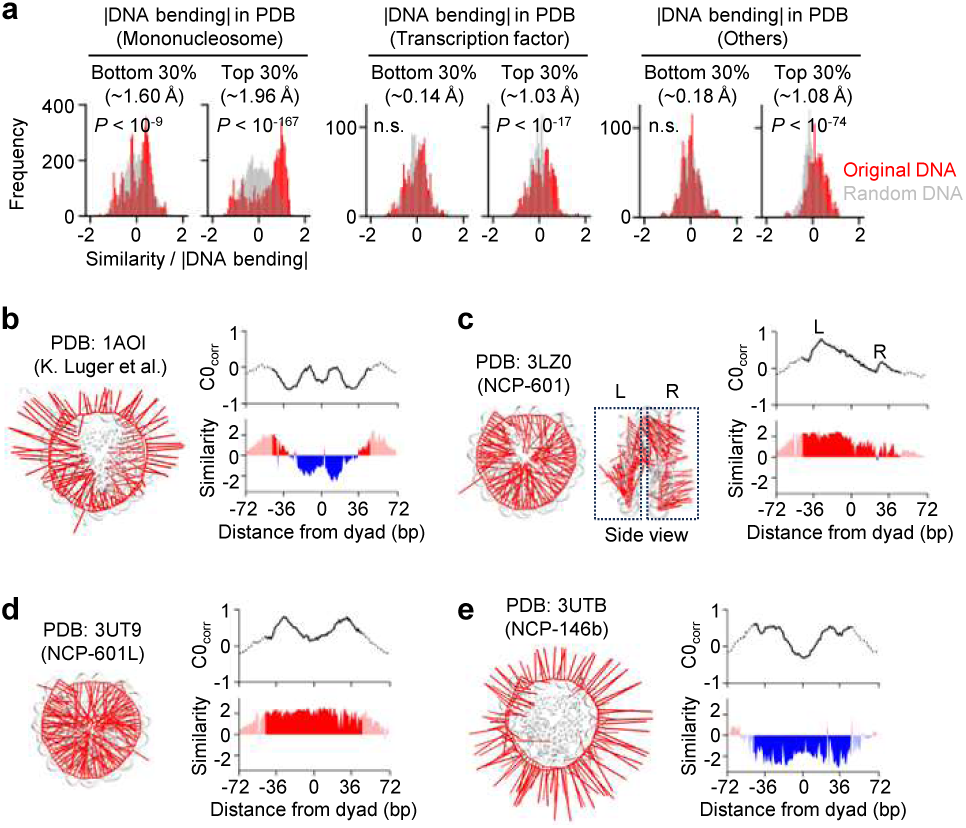
Selection of DNA mechanics in experimentally determined structures. **a**, Selection of DNA mechanics in different molecular types. *P*-values by the paired t-tests between randomized and original DNA sequences are shown. **b**, **c**, **d**, **e**, DNA mechanical properties of complexes shown in PDB IDs of 1AOI, 3LZ0, 3UT9, and 3UTB, respectively. 3-dimensional DNA bending is overlaid with PDB structures (left), and the corresponding cyclizability and similarity scores are plotted (right).

**Supplementary Fig. 7.**
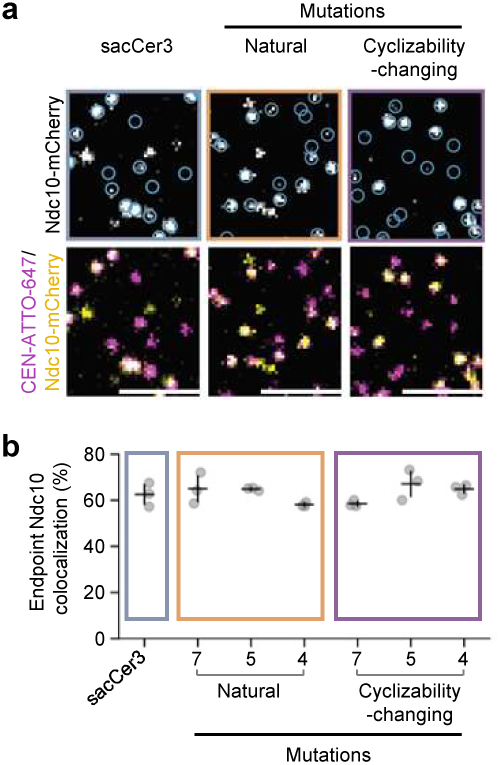
Colocalization of centromere DNA and Ndc10 in inner kinetochore. **a**, Example images of total internal reflection fluorescent microscopy endpoint colocalization assays of visualized Ndc10-mCherry on sacCer3 CDEII DNA (left), CDEII with natural mutations (middle) or cyclizability-changing mutations (right), with colocalization shown in relation to identified DNA in blue circles. Bottom panels show respective overlays of DNA channel (magenta) with Ndc10-mCherry (yellow). Scale bars: 3 μm. **b**, Graph indicates quantification of endpoint colocalization of Ndc10 on sacCer3 CDEII DNA (left), CDEII with natural mutations (middle) or cyclizability-changing mutations (left). Points indicate individual experiments (n=3) where ∼1,000 DNA molecules were identified per replicate.

**Supplementary Fig. 8.**
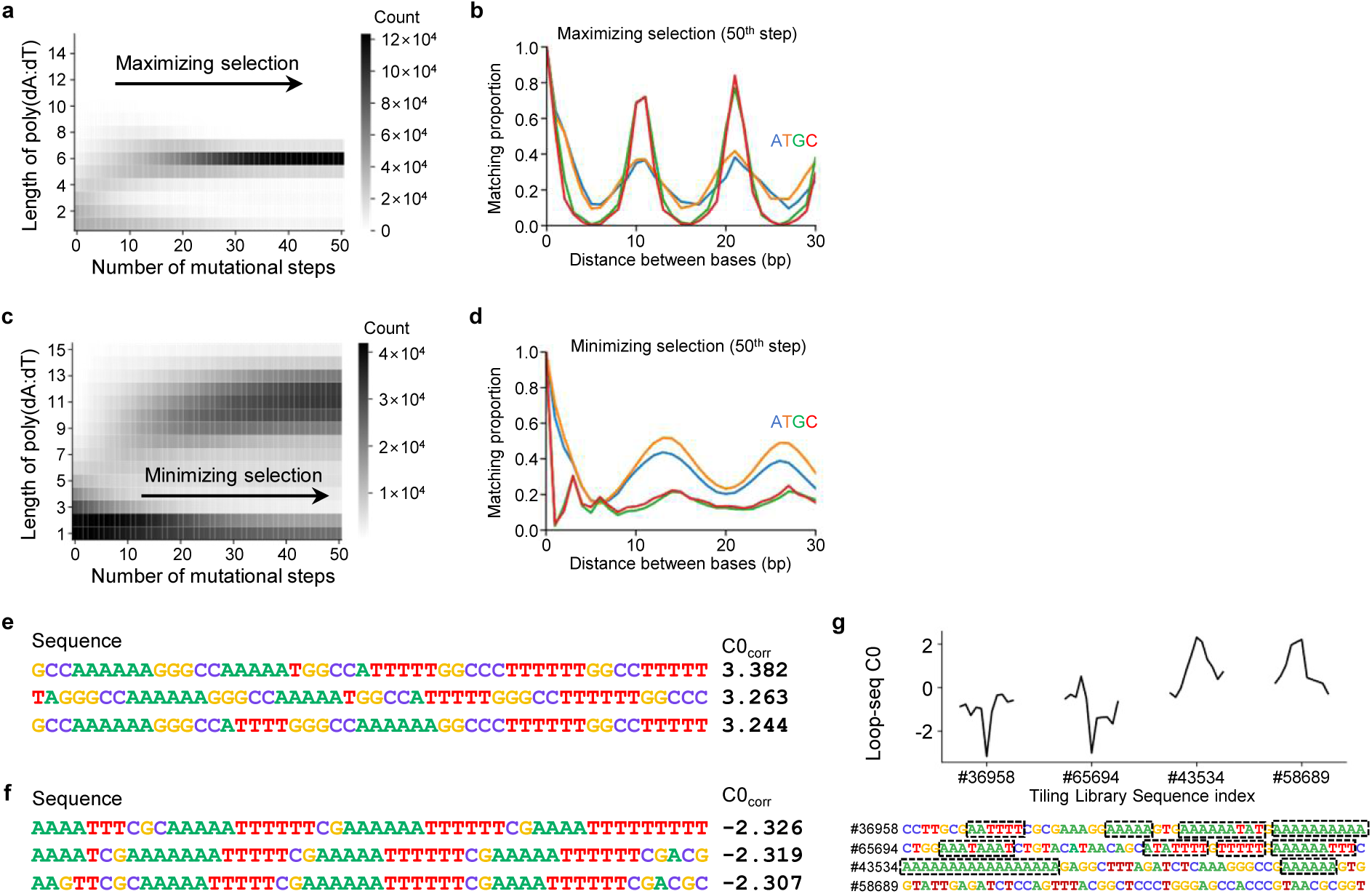
Accumulation of poly(dA:dT) tracts during *in silico* evolution. **a**, The number of bases that belong to a poly(dA:dT), defined as runs of dAs or dTs of the indicated length found in the pool of sequences under maximizing selection (Fig. 4c). **b**, Proportion of matching bases of the same type at each distance after 50 maximizing steps. For example, the sequence 5’-NN<u>A</u>NN<u>A</u>NN-3’ is used to count adenine (A) matches at a distance of 3 bases. **c**, **d**, Repeat of **a** and **b** but for minimizing selection, respectively. **e**, Three sequences with the highest predicted C0_corr_ found after 50 steps of maximizing selection. **f**, Repeat of **e** but for the lowest predicted C0_corr_. **g**, Two sequences with the lowest and two sequences with the highest measured C0 in the Tiling library. The measured C0 of adjacent sequences was plotted together. Poly(dA:dT) tracts longer than 5 bp are highlighted in the dashed boxes. **a**, **c**, **g**, Any continuous fragment of dA, dT, or their mixtures, such as 5’-ATTATAT-3’, is considered a poly(dA:dT) tract.

**Supplementary Fig. 9.**
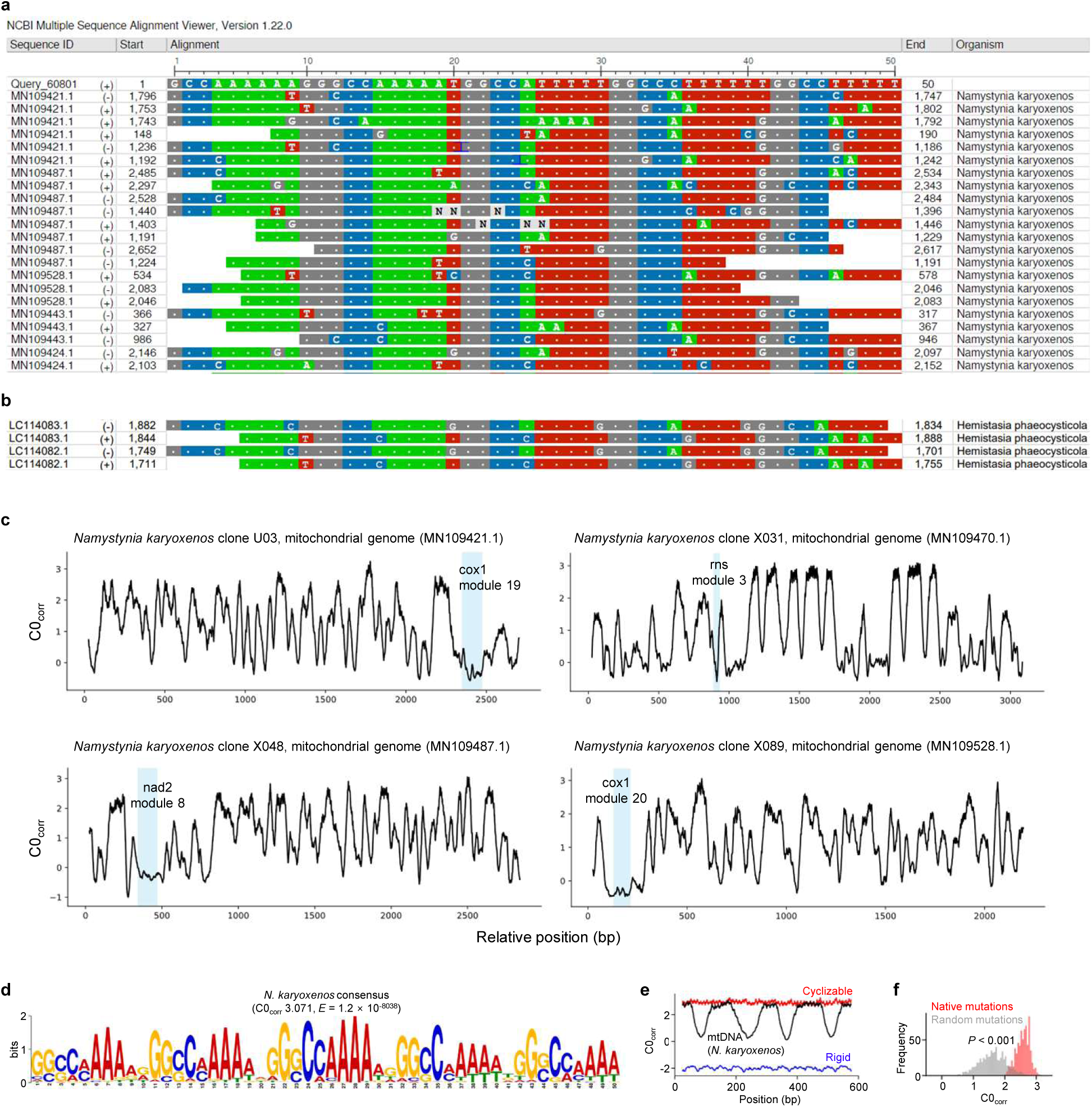
Extreme DNA mechanics in Hemistasiidae mitochondrial genomes. **a**, Top hits from the BLASTn alignment results using the 50 bp DNA with the highest C0_corr_ found after the maximizing selection (the first sequence of Supplementary Fig 8e, Methods). **b**, Mitochondrial genomes of *H. phaeocysticola* found in the same BLASTn search in **a**. **c**, C0_corr_ of the four regions of mitochondrial genomes of *N. karyoxenos* containing unusually high C0_corr_. Coding regions are highlighted in the blue shaded areas. **d**, The repetitive 50 bp motif found in the mitochondrial genomes of *N. karyoxenos*, based on MEME analysis^79^. **e**, C0_corr_ vs position for the three 600 bp DNA used in AFM images in Fig. 4e. **f**, Natural mutations in the mitochondrial repeat motifs rarely show C0_corr_ below 2, but random mutations do. The Hamming distance for the native and random mutations was preserved (Methods).

**Supplementary Fig. 10.**
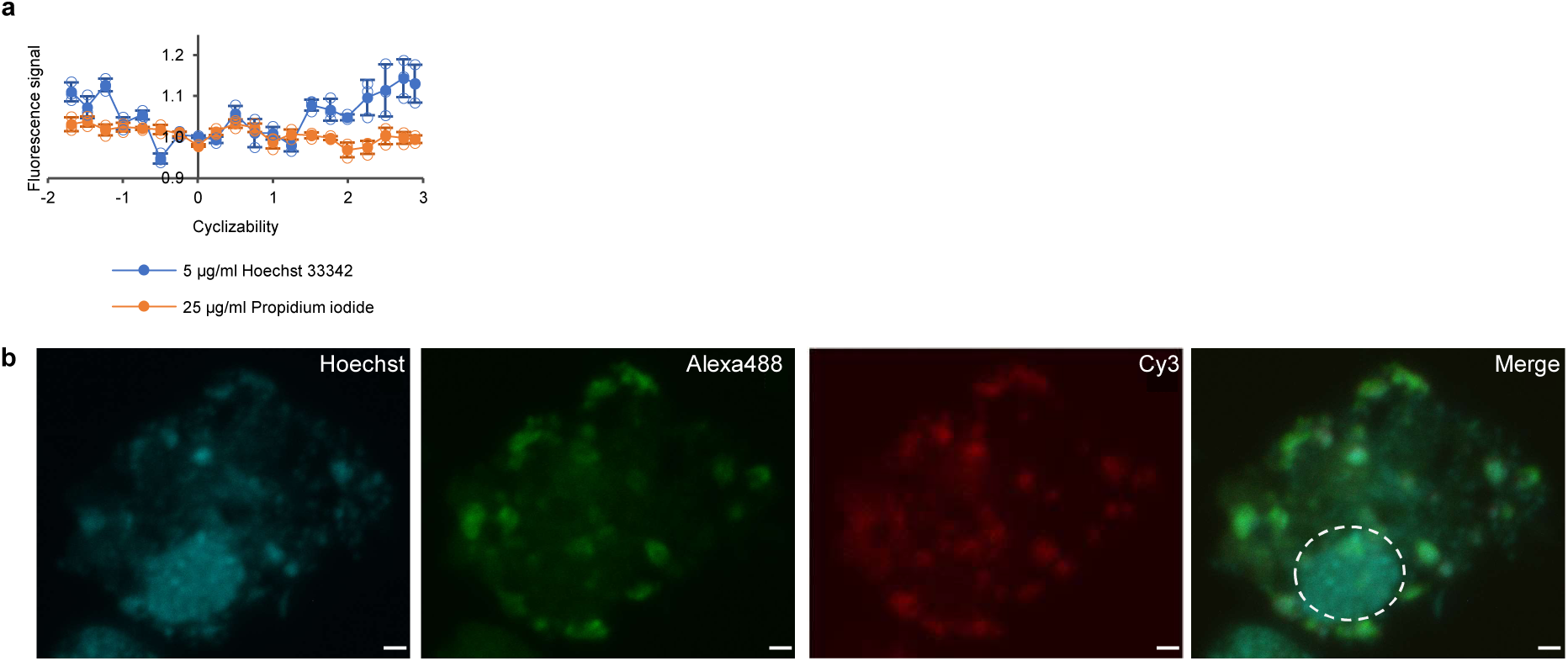
Micrographs of nucleus and mitochondria of *N. karyoxenos*. **a**, Relative *in vitro* Hoechst 33342 and PI fluorescence intensity when mixed with dsDNA with a range of cyclizabilities. Fluorescence signal intensity was normalized to the mean of three samples with close zero cyclizability. Open circles are data points from individual replicates (n = 3), solid circles represent the mean, error bars represent the standard deviation. **b**, Overlay of z-stack images from confocal microscopy showing combined immunofluorescence (IF, Alexa488) and fluorescence *in situ* hybridization (FISH, Cy3) analysis. IF assay was performed using an antibody against the β chain of mitochondrial ATP synthase, FISH using a DNA-Cy3 probe which labelled circularized mtDNA regions. Dashed lines encircle the nucleus. Scale bar: 1 µm.

**Supplementary Fig. 11.**
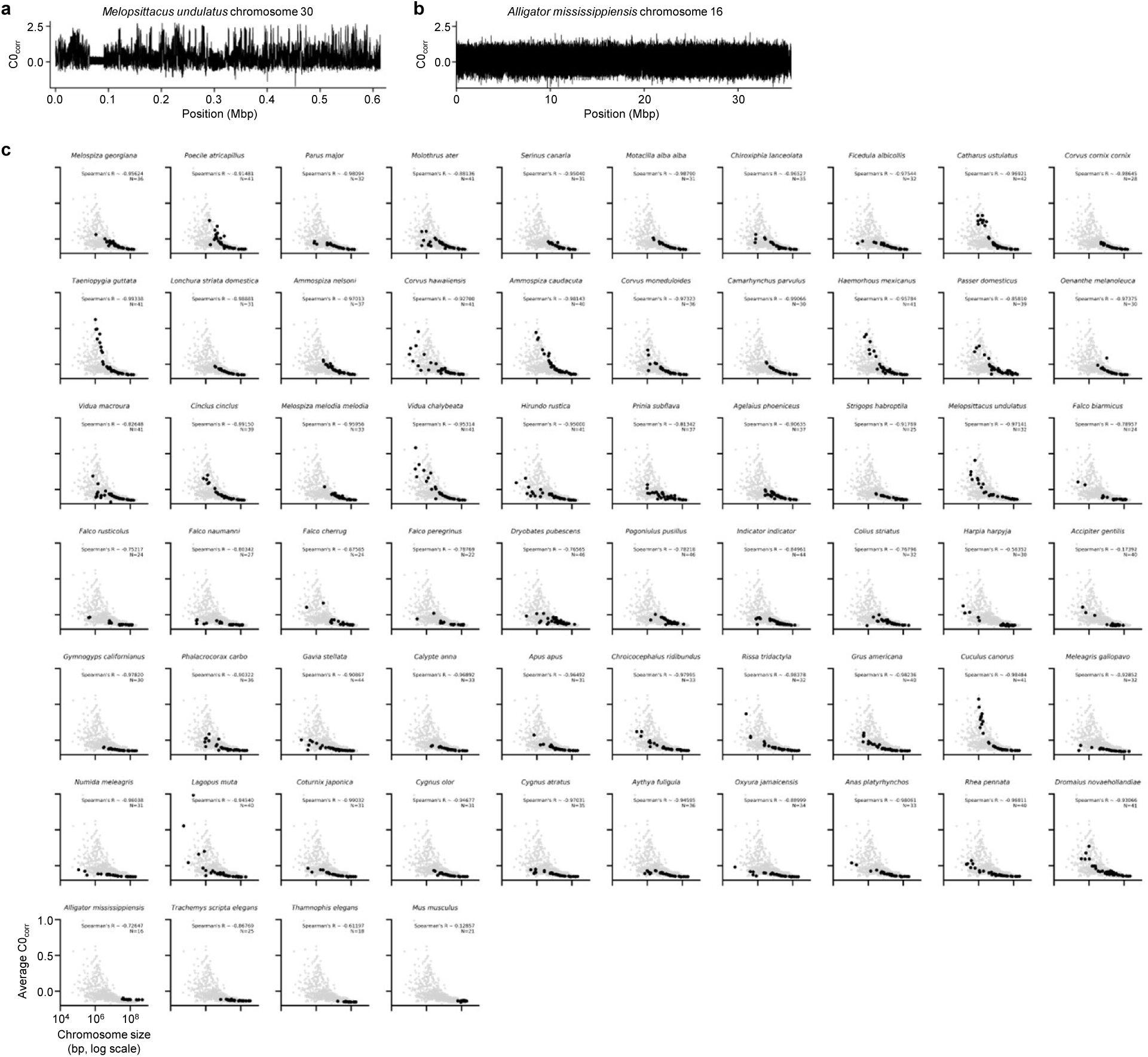
Cyclizability of two example chromosomes. **a**, Cyclizability along chromosome 30 of *M. undulatus*. **b**, Cyclizability along chromosome 16 of *A. mississippiensis*. **c**, Chromosome size vs average cyclizability per chromosome for 60 bird species and 4 outgroups. Gray dots indicate the distribution of all chromosomes collected from 60 bird species.

