## Supplementary materials for "Probing mechanical selection in diverse eukaryotic genomes through accurate prediction of 3D DNA mechanics"

### Supplementary information

#### Table of contents

Supplementary Table 1. Nucleotide information

Supplementary Note 1. Adapter-dependent cyclizability

Supplementary Note 2. 95% CI of measured cyclizability

Supplementary Note 3. Details for training models to predict adapter-dependent cyclizability

Supplementary Note 4. Adapter-corrected cyclizability

Supplementary Note 5. Bending of $(dA)_n$ in 3-dimensional space

Supplementary Note 6. Quantifying mechanical selection

Supplementary Note 7. Details for quantifying mechanical selection in centromeres

Supplementary Note 8. Heuristics to accelerate cyclizability predictions for long DNA sequences

#### Supplementary Table 1. Nucleotide information

DNA constructs used for smFRET DNA cyclization assay (Fig. 4d, g). T highlighted with yellow was linked to a biotin.

| | |
|---|---|
| High $C0_{corr}$ DNA | 5′Cy3CAGAATCCGTCGAAGAGCGGCCAAAAAAGGCCGTTTTTGGGCCATTTTTGGCCCTAAAAAGGGCTTTTGCTCTTCG3′<br><br>3′GCTTCTCGCCGGTTTTTTCCGGCAAAAACCCGGTAAAAACCGGGATTTT**T**CCCGAAAACGAGAAGCGTCTTAGGCACy55′ |
| Low $C0_{corr}$ DNA | 5′Cy3CAGAATCCGTCGAAGAGCAAAAAAAATTTTTCGAAAAATTTTTCGAAAAAATTTTTTCGAAAATTTTTGCTCTTCG3′<br><br>3′GCTTCTCGTTTTTTTTAAAAAGCTTTTTAAAAAGCTTTTTTAAAAAAGC**T**TTTAAAAACGAGAAGCGTCTTAGGCACy55′ |

DNA constructs used for AFM imaging (Fig. 4e, f).

| | |
|---|---|
| mtDNA<br><br>(*N. karyoxenos*) | 5′GGCCCATTTTGGGCCCATTTTGGGCCCTTTTTGGGCCCATTTTGGGCCCTTTTGGGCCCTTTTTTGGCCCTTTTCTCCGGGGGTCCCTG<br>TCCTGTTGGGGTTTTTGGCCCTTTTTGGCCCATTTTTGGGCCATTTTGGCCCAAAATGGCCCAAAAAAGGGCCAAAAATGGCCCAAAATAG<br>GCCATTTTGGGCCATTTTGGCCCCTTGGGGCCCAAAATAGTAGAATGTAACAGTATAGGCCCTATGTAATTGGGCCTAGGTGGCCCAGGGG<br>CCCAAAATGGGCCCAAAAGGGCCCAAAATGGGCCCTTTTTGGGCCCTTTTTGGGCCCTTTTTGGCCCCTTTTTGGCCCTTTTCTCTGGGGG<br>TCCCTGTGCTGTTGGGGTTTTGGGCCCTTTTTGGCCCATTTTAGGCCCAAAATGGCCCAAAATGGCCCAAAAATGGCCCTTTTTGGGCCCT<br>TTTTGGGCCCTTTTTGGGCCCTTTTTGGGCCCTTTTTGGCCCTTTTCCCTGGGGGTCCCTGTCCTGTTGGGGTTTTGGGCCCATTTTGGGC<br>CCAAAATGGGCCCTTTTTGGCCCAAAATGGGCCATTTTGGGCCCTATTTTGGCCTT 3′ |
| Cyclizable | 5′TTTAGGCCCATTTTTGGCCAATTTTGGGCCAAAAAAGGGCTAAAAAAGGCCAATTTTGGCCCATTTTGGGCCATTTTTGGGCCATTTTT<br>GGCCCTAAAAAGGCCATTTTTGGCCGTTTTTGGGCCAAAAAAGGGCCAAAAATGGCCTTTTTAGGGCCAAAAAAGGGCCAAAAATGGCCAT<br>TTTTGGGCCAAAAATGGCCCTTTTTGGGCCATTTTTGGGCCAAAAAAGGCCATTTTTGGGCCTTTTTTGCGCCAAAAAAGGCCATTTTTGG<br>GCCATTTTTGGCCATTTTTGGGCCTAAAAAGGGCCAAAAAAGGGCATTTTTGGGCCAAAAAAGGCCCAAAAATGGCTTTTTTAGGCCATTT<br>TTGGCCTATTTTGGGCCATTTTTGGGCATTTTTTGGCCTTTTTGGGCCCTTTTTGGGCCATTTTTGGCCCAAAAATGGCCATTTTTGCGCC<br>AAAAAAGGGCCAAAAATGGCCATTTTTGGCCCAAAAATGGCCTAAAAGGGCCATTTTTGGGCCAAAAAAGGCCAAAAAAGGGCCTTTTTTG<br>GCCTAAAAAGGCCCAAAAATGGCCATTTTTGGCCCATTTTTGGCCCAAAAATGGCC 3′ |
| Rigid | 5′AAAAAAAATTTTTCGAAAAAATTTTTTCGAAAAAATTTTTTGAAAATTTTTGGTCGCGAAAAATTTTTCGCGAAAAATTTTTCGAAAAT<br>TTTTTCGACGAAAAAATTTGTCGCAAAAATTTTTCGAAAAATTTTTGATCGAAAATTTTTTCGAGTCGAAAATTTTCGAAAAAATTTTTCG<br>ACAAAAATTTTTCGAATTTTTTTTTTCGCGAAAAATTTTTTCGAAAAAATTTTTTCGAAAATTTTTCGACGAAAAAATTTTCGAAAAATTTT<br>TTCGAGTCAAAATTTTCGAAATTTTTTTTTCGAAAAATTTTTGACGAAAAAATTTTTTCAAAAATTTTTTTCGAATTTTTTTTCGAAAAAT<br>TTTTCGCGAAAAATTTTTTCGAAAAATTTTTTCGAACTAAAATTTTCGAAAAATTTTTCGAGTCAAAATTTTTGAAAATTTTTTCGACAAA<br>ATTTTTTCGCGAAAAATTTTTCGAAAAAATTTTTTTCGAAAATTTTTCGTGATGAAAAATTTTTGAAAAAATTTTTTCGAAAAATTTTTTC<br>GAATTTAAAATTTTGAAAAAAAATTTTTTCGAAAAAATTTTTTCGACGTCCGATTTT 3′ |

| | |
|---|---|
| Natural mutations | 5’AATATTTCATTTTATTATATTTTAAAAAAAAGTAAAAAATAAAAAGTAGCTTATTTTCAAAAAATAAAATCTACAATATTAG 3’ |
| | 5’AATATTTGATTTTATTATATTTTTAAAAAAGGGAAAAAATAAAAAGTAGTCTATTTTTAAAAAGTAAAATTTAAAATATTAG 3’ |
| | 5’GATATTTGATTTTATTATATTTTTAAAAAGAGTAAAAAATAAAAAGTGGTTTATTTTCAAAAAATAAAATTTAAAACATTAG 3’ |
| Cyclizability changing mutations | 5’TATATCTGATTTTATTATATTTTTAAACAAAGTAAAACATAAAAAGTAGTTTAATTTTATAAAATAAGATTTAAAATATTAG 3’ |
| | 5’GATATTTAATTTGATTATATTGTTAAAAAAAGTAAAAAATAAAAAGTAGTTTATGTTTAAAAAATAACATTTAAAATATTAG 3’ |
| | 5’GATATCTGATTTTACTATATTTTTAAAAAAAGTAAAGAATAAAAAGTAGTTTATTTTTAAAAAATACAATTTAAAATATTAG 3’ |

CDEII DNA constructs used for centromere protein colocalization assay (Fig. 3e-i, Supplementary Fig. 7).

DNA constructs used for Hoechst 33342 affinity test (Supplementary Fig. 10a). DNA sequences are sorted by $C0_{corr}$ (from

| |
|---|
| 5’GCCGGGCGCAAATTTTGACGTCGTCGACCGACGTCAAAATTTTTCGTCGAAATTTTTTTCGCGAAATTTTTTTCGCCCAAATTTGACGACGCGCCCGCCC3’ |
| 5’CGCCGGCGGCGGAAGTCGAAATTTTCGACACGGAAAATTTCGGACTCGGAAAATTTTTCGACCGAGTCGTGACGCGAAATTTTTGGCCGACATTTTTCCC3’ |
| 5’GACAATTTCGGATCGTCGCGACGACTCGGCGACTCAACTAAAATTTTTTCGAACGGTGACGCGAACTTCGACTCGCGAAATTTTTCCGAACGCACGTGGT3’ |
| 5’GGAGTCTGACGACTCGACCGAGTGTTCGACGATTTCCGACTTTTGAAAAAACGCGGTCAGCGAATTTTGGTCACGGAGTCCTTCGCGATTTTCCGATCTT3’ |
| 5’CTTGACGAAATGTCAAGCGAGGGCGTCCTTGTCATCAAGTGCATTTTATTTTTCCACGCGTCGAAGGAATTACGACTTCGACCTCACACTCGCGAACGCC3’ |
| 5’CAGAACTTATCCTGATGGGCGGACGTTAGTAAGACTCATCCTTGTCTTGATCGACCGTCCACATTAAGTTACCAGTCTTGCTCGTCTCACACGAGGCGTG3’ |
| 5’TTTTCCGCCGCACCTCTGCATTATTGTACTGTGTCTTAATACCGGCGGTTGGTCTAGTACGCTGCACTTGAATACACAGCGCTGGTCTTACCAACTAGCC3’ |
| 5’ACATTTAGCGCCTGTGCCCCACATACGCAGGCGTCCTATATTGTACGGATATCTTTACCTCTATAGCTTACCATGTTAGGGGCCGCCCCCAGGATATAGT3’ |
| 5’TTCGTGGGTGGATAGTACCTCCTTTCAGGGATTAGAGCCTAATAGGTGAACAGTAACGGCGGTTCTCCCTCATTCAGAGGCATGTAAGGGAAGGCTCAGT3’ |
| 5’AAGGACCCGATAACAGGGTGCAAGACGCGTATATCTACAGCATCCGGGCTATATTGAACGCTCAGAACTGTACAAACCCCTCAACGGAAGGCATGGAGTA3’ |
| 5’ACCAGCGTGTCCAGACCCGATCAGACCCGCGTTTTGCACTCTTCTTATATAAAAACACCTCGAGAAGCATAGTTAGAGGGAGCCCAGGCTCAATACAGAC3’ |
| 5’TAGCCATGCAAGGACCATTTGGGGACTTCAGAGGGGGTACTTTCGTCCCAATTAAAGGCCTAGAGGATGCCAATTGGACTCTCGTGAGGTACAATTCACC3’ |
| 5’GTGGCACTTTTTGGCTCTATTTAAGCACTCCTGTGCTACCTAACAGGCGAATTTACGGGTAAAAACGGGTGCGTTAGGCATCCCGAGCCGCTCGTTTTAC3’ |
| 5’CGCGCCATATGGCTGTTTTTGCGGCGAAATTCACCCCTAATAGGAGCTGTAAAGCCGAAAATAGGGCGGAAATCGCTCCGTAAAGGCAATAGAGCAGCTT3’ |
| 5’GGGCTCTTTCGAGGCCTCTTTTGGGGTCCAAAACGGCGCCAAACGCATAAAAAGCGTGTAATTTAGGGCATTTGACGCTATTTCACGCCTTCTGGACCAA3’ |
| 5’AGGCCTAAAATAGCCCCCAGAATGGCTCCAGAGGGACCGTTTTGGGCCGAAAAAAGGCCTCTAAATAGGCCGAAACCCGCGAATTTCGCGTCTATTTTAA3’ |
| 5’AAAAAAGTACAAAAAACGGCCCTTAGGAGCCTGTTTTAGGGGCTTTTTGGGCCAAAATACGCCCGAAAAGGCGCTTAGGAGGCCTCTGAAAGGGCCTGAG3’ |
| 5’CAGGGCGAAAAAAGGCCAAAAGAGCCCTTTTTGGCCCATAAAATGGCCTATTTTGGCCCAAAAAGGCCAATTTTAGCCCCCCAGAGGGCGAATTTTGCGC3’ |
| 5’CTTAGGGCGTCCCAGAGGCCCCAAAAAGGCCATTTTTGGGCCAAAAAACGCCAAAAAAGGGCGAAAAAGGGCCGAAAAAGGCCTAATTTGGCCCGAAATT3’ |
| 5’AAAAGGGCTCAAAAAAGGCCCAAAAAGGGCCAAAAAAGGCCCAAAAAAGGCCATTTTTGGCCGTTTTTGGGCCTCTAAAGGGCCCCAGAGGCCCTCGATA3’ |

lowest to highest).

**Supplementary Note 1. Adapter-dependent cyclizability**

Adapter-dependent cyclizability, originally described in Basu et al. (Nature 2021)[1] is a quantity of a DNA sequence (50 bp in most published studies) measured by loop-seq. C26, C29, and C31 values are directly measured using loop-seq, and C0 is inferred from C26, C29, and C31 using a trigonometric formula. In this section, we outline the definitions of cyclizability.

For loop-seq, DNA molecules are prepared with variable 50 bp sequences in the middle and 25 bp adapters with 10 bp ssDNA overhangs at both ends. The 25 bp adapter downstream of the central 50 bp is bound to a bead surface via a biotin tether (Supplementary Fig. 1). A DNA library sample composed of such DNA molecules is split into two halves and treated with 1 M NaCl to induce DNA looping. For the first half, unlooped DNA molecules remaining after the 1 M NaCl treatment are digested by the exonuclease RecBCD. RecBCD digestion is omitted for the second half (no-digestion control). Each half is then amplified and sequenced (Fig. 1a).

In the DNA sample digested by RecBCD, we define $N_s$ as the total read counts and $n_s$ as the read count of a specific sequence whose cyclizability is to be measured. $N_c$ and $n_c$ are equivalent quantities for the no-digestion control. Cyclizability is defined as follows.

$$C = \log\big((n_s/N_s)/(n_c/N_c)\big) = \log(n_s/n_c) + \log(N_c/N_s)$$

Here, $\log(N_c/N_s)$ is a constant equally applied to every sequence in the library and is occasionally omitted from the definition. Since this constant varies among experiments, cyclizability measured by different loop-seq experiments can be compared up to a constant offset.

The biotin tether and bead surface restrict DNA looping in 3-dimensional space such that cyclization is favored for DNA sequences that have intrinsic bending direction away from the biotin tether. In most published studies, three different positions of biotin tether were used, which are 26, 29, or 31 nucleotides (counting from the 5' end of the downstream ssDNA overhang) corresponding to C26, C29, and C31. Thus, C26, C29, and C31 measure the propensity of DNA bending toward three different directions. We can fit different cyclizability values into a trigonometric formula as follows.

$$C(n) = C0 + A\sin(kn + \varphi), \text{ where } n = 26, 29, 31 \text{ and } k = \frac{2\pi}{10.3}\, bp^{-1}$$

C0 is an intrinsic cyclizability of a sequence, independent of the rotational phasing set by biotin tethers[1].

#### Supplementary Note 2. 95% CI of measured cyclizability

We defined $N$, $n$, $r$, and $C$ as follows.

$N$ : The total number of aligned reads of DNA molecules in the pool.

$n$ : The number of sequencing reads aligned to the target DNA sequence. $N$ is the sum of all reads of different DNA sequences ($N = \sum n$).

$r$ : The ratio of molar concentrations of the target DNA molecule and total DNA in solution. $r$ is measured as $\frac{n}{N}$.

$C$: Cyclizability. The lower cases c and s stand for control (sequenced without digestion) and sample (sequenced after digestion of unlooped DNA molecules). Cyclizability is defined as $\log(r_s/r_c)$, which is measured using the read counts as $\log\big((n_s/N_s)/(n_c/N_c)\big)$. We refer to $\log(r_s/r_c)$ as ideal cyclizability and $\log\big((n_s/N_s)/(n_c/N_c)\big)$ as measured cyclizability.

We obtained a 95% CI for cyclizability to quantify the inaccuracy in loop-seq measured cyclizability using Bayesian inference. Assuming the Poisson distribution, we get

$$P(n|r) \; = \; \frac{1}{n!}(Nr)^n e^{-Nr}.$$

The prior $P(r)$ was assumed to be proportional to $\frac{1}{\sqrt{r}}$, a non-informative Jeffreys prior of the Poisson distribution. We may adopt different priors, such as a uniform prior, which does not make a significant difference to the results. In this study, we used the Jeffreys prior by setting $P(r) \sim \frac{1}{\sqrt{r}}$.

$$P(r|n) \sim P(n|r)P(r) \sim \frac{1}{n!\,\sqrt{r}}(Nr)^n e^{-N} \quad \sim \; r^{n-\frac{1}{2}}e^{-Nr}$$

We define $f(r) \; \coloneqq \; r^{n-\frac{1}{2}}e^{-Nr}$ by leaving $N$ and $n$ as constants.

$$P(C|n_c,\, n_s, N_c, N_s) \; \sim \; \int P(r_c|n_c, N_c)P(r_s|n_s, N_s) \sim \; \int f(r_c)f(r_s)$$

We define variables $R$, $\theta$ that satisfy $r_c \; = \; R\cos\theta$ and $r_s \; = \; R\sin\theta$ ($R > 0, 0 \leq \; \theta < 2\pi$).

We defined $g(R,\, \theta)$ as follows.

$$g(R,\, \theta) \; \coloneqq \; f(r_c)f(r_s) \; = \; r_c^{\,n_c-\frac{1}{2}}e^{-N_c r_c}\, r_s^{\,n_s-\frac{1}{2}}e^{-N_s r_s}$$

Then we get $g(R,\ \theta) = (\cos\theta)^{n_c-\frac{1}{2}}(\sin\theta)^{n_s-\frac{1}{2}}R^{n_c+n_s-1}e^{-(N_c\cos\theta+N_s\sin\theta)R}$.

By replacements $a = n_c + n_s - 1$, $b = N_c\cos\theta + N_s\sin\theta$, and $l = \min\left(\sec\theta,\ \sec\left(\frac{\pi}{2}-\theta\right)\right)$ we got

$$P(C|n_c,\ n_s,\ N_c,\ N_s) := \int_R g(x,\ \theta)dx = \int_0^l (\cos\theta)^{n_c-\frac{1}{2}}(\sin\theta)^{n_s-\frac{1}{2}}x^a e^{-bx}dx$$

$$= (\cos\theta)^{n_c-\frac{1}{2}}(\sin\theta)^{n_s-\frac{1}{2}}\left[b^{-(a+1)}\Gamma(a+1,\ bx)\right]_0^l$$

$$\approx (\cos\theta)^{n_c-\frac{1}{2}}(\sin\theta)^{n_s-\frac{1}{2}}b^{-(a+1)}\ .$$

Then, the following relations hold.

$$C = \log(r_c/r_s) = \log(\tan\theta),\ e^C = \tan\theta,\ \sqrt{e^{2C}+1} = \sec\theta,$$

$$\cos\theta = \frac{1}{\sqrt{e^{2C}+1}},\ \text{and}\ \sin\theta = \frac{e^C}{\sqrt{e^{2C}+1}}.$$

Using the relations above, we obtained,

$$P(C|n_c,\ n_s,\ N_c,\ N_s) \approx (e^{2C}+1)^{\frac{1}{2}}e^{C\left(n_s-\frac{1}{2}\right)}(N_c + N_s e^C)^{-(n_c+n_s)},$$

which is the probability distribution of the true cyclizability of a 50 bp DNA sequence.

The 95% CI of cyclizability is defined as $C_{lower} < C < C_{upper}$, where $P(C \le C_{lower}\mid n_c, n_s, N_c, N_s) = 0.025$ and $P(C \ge C_{upper}\mid n_c, n_s, N_c, N_s) = 0.025$. The probability $P(C|n_c,\ n_s,\ N_c,\ N_s)$ was computed for every $C = 0.01t$, where *t* is an integer. Cyclizability with a probability lower than $e^{-8}$ was ignored. We obtained the highest $t_1$ that holds $\sum_{t=t_1}^{\infty}\frac{P(0.01t)}{I} > 0.025$, and the lowest $t_2$ that holds $\sum_{t=-\infty}^{t_2}\frac{P(0.01t)}{I} > 0.025$, where $I = \sum_{t=-\infty}^{\infty}P(0.01t)$. The 95% CI of cyclizability was determined as $\left(0.01\left(t_2-\frac{1}{2}\right),\ 0.01\left(t_1+\frac{1}{2}\right)\right)$.

Alternatively, we used frequentist statistics to estimate the uncertainty in the measured cyclizability. We first defined $\hat{r}$, the observed proportion of the target sequence in the pool of DNA. $\widehat{r_c}$ and $\widehat{r_s}$ stands for $\hat{r}$ of control and sample, respectively. Assuming the normal distribution of $\hat{r}$, we get

$$\hat{r} \sim N\left(r, \sqrt{\frac{r(1-r)}{N}}\right) \approx N\left(r, \sqrt{\frac{r}{N}}\right)$$

We approximated $1 - r \approx 1$ as $r$ does not exceed 0.001 in most cases. Accordingly,

$$R := \frac{\hat{r}}{r} \sim N\left(1, \frac{1}{\sqrt{Nr}}\right) \approx N\left(1, \frac{1}{\sqrt{n}}\right)$$

We defined the observed cyclizability, $\hat{C} := \widehat{r_s}/\widehat{r_c}$.

$$\frac{e^{\hat{C}}}{e^{C}} = \frac{\left(\frac{\widehat{r_s}}{\widehat{r_c}}\right)}{\left(\frac{r_s}{r_c}\right)} = \frac{\left(\frac{\widehat{r_s}}{r_s}\right)}{\left(\frac{\widehat{r_c}}{r_c}\right)} = \frac{R_s}{R_c}$$

By adopting $V\left(\frac{1}{R}\right) \approx \frac{V(R)}{[E(R)]^4} = V(R)$ and assuming the independence of $R_c$ and $R_s$, we derived

$$V\left(\frac{e^{\hat{C}}}{e^{C}}\right) = V\left(\frac{R_s}{R_c}\right) \approx [E(R_s)]^2 V(R_s) + \left[E\left(\frac{1}{R_c}\right)\right]^2 V\left(\frac{1}{R_c}\right) + V(R_s)V\left(\frac{1}{R_c}\right)$$

$$\approx V(R_s) + V(R_c) + V(R_s)V(R_c) \approx \frac{1}{\sqrt{n_c}} + \frac{1}{\sqrt{n_s}} + \frac{1}{\sqrt{n_c n_s}}$$

Thus, we get $\frac{e^{\hat{C}}}{e^{C}} \sim N\left(1, \frac{1}{\sqrt{n_c}} + \frac{1}{\sqrt{n_s}} + \frac{1}{\sqrt{n_c n_s}}\right)$. The variance $\frac{1}{\sqrt{n_c}} + \frac{1}{\sqrt{n_s}} + \frac{1}{\sqrt{n_c n_s}}$ represents the uncertainty of the measured cyclizability. The variance correlates strongly with 95% CI that we previously obtained using the Bayesian inference (Spearman's $R$ = 0.997, Fig. S1). The variance, or an 'uncertainty score' (approximated by frequentist statistics), is calculated in a constant computational complexity, while the Bayesian inference requires computations with a polynomial complexity. Thus, we used the uncertainty score instead of 95% CI when precise intervals of 95% CI were not required.

**Supplementary Note 3. Details for training models to predict adapter-dependent cyclizability**

In the previous study[1], each read count was increased by 1 before obtaining cyclizability, because cycilzability is not defined for sequences with zero reads. However, in this study, sequences used in model training were first screened by the uncertainty score to remove sequences with low reads, thus, it is possible to calculate cyclizability without adding an extra constant to read counts.

During model training, we removed sequences that included 7-mer Nt.BspQ1 recognition motifs (Methods). The absence of some 7-mer motifs in the dataset may lower the prediction quality on the sequences including such 7-mer motifs. We grouped the previous datasets based on whether a sequence includes an arbitrarily chosen pair of reverse complementary 7-mer motifs, 5'-CGAGAAG-3' and 5'-CTTCTCG-3'. Sequences without the motifs were collected for the training dataset, and the remainders were used for the validation of the model. The model structures and training scheme were not modified. The models have shown comparable prediction accuracy for the sequences including the 7-mer motifs, 5'-CGAGAAG-3' or 5'-CTTCTCG-3' (Fig. S2). Still, any analysis of the sequences containing the recognition motifs of Nt.BspQ1 enzyme should be aware of the possible inaccuracy of the predictions.

To see if read counts affect prediction accuracy, we used repeated measurements of C26 of the Cerevisiae Nucleosome library[1]. For each library member, among the two measured values of C26, one with a with a smaller 95% CI width agreed better with predicted C26 (Pearson's R = 0.906, Supplementary Fig. 4b) than the other C26 with a larger 95% CI width (Pearson's R = 0.860, Supplementary Fig. 4b).

As a further test, we grouped sequences in the Random library into 8 subsets based on their 95% CI of C26. Prediction was more accurate on subsets with the smallest 95% CI of measured C26 width, and a similar pattern was observed in the ChrV library (Supplementary Fig. 4c).

#### Supplementary Note 4. Adapter-corrected cyclizability

Biotin tethering of DNA molecules in loop-seq experiments restricts DNA looping at certain geometry. When biotin is attached to the 26th base from the right end of the DNA (Supplementary Fig. 1), the DNA cannot bend at the 26th base toward the bead (Fig. 2a). Cyclizability measured under this setting is named C26, and a similar rule applies to C29 and C31. Intrinsic cyclizability, C0, is inferred from three cyclzabilities, C26, C29, and C31, after fitting them into a formula.

$$C(n) = C0 + A\cos\left(\frac{60.5 - n}{10.3} * 2\pi - \frac{2}{3}\pi - \varphi\right), \qquad where\ n = 26, 29, 31 \tag{1}$$

Assigning C26, C29, and C31 to the formula above determines the intrinsic cyclizability, amplitude $A$ and the phase $\varphi$. Ideally, the amplitude is the propensity of DNA looping in a certain direction described by the phase. The terms $\frac{60.5-n}{10.3} * 2\pi$ and $-\frac{2}{3}\pi$ were included so that $\varphi = 0$ implies the base in the middle of the variable sequence (60th and 61st bases from the 5' end of a ssDNA overhang, or 25th and 26th bases in the variable 50 bp DNA, Supplementary Fig. 1) bends toward the minor groove.

According to this definition, C0 is determined by the synergic effect of the variable sequence and the adapter sequences, as C26, C29, and C31 measurements include the adapter sequence in the experimental setting. As a result, the reverse complementary sequences have C0 that is different from that of the original sequences (Fig. 1e). Here, we suggest a method to reduce the effect of adapter sequences on cyclizability.

##### Revisiting the formula

For a sufficiently long DNA that prefers bending to a single direction, the phase $\varphi$ changes by $\frac{2\pi}{10.3}$ for every base pair due to the helical structure of DNA. Accordingly, C26, C29, and C31 oscillate across the region (Supplementary Fig. 5a, left). The oscillations are contained within a pair of hypothetical envelopes (Supplementary Fig. 5a, left). The envelopes were determined for each cyclizability, $C(n)$, where $n$ is 26, 29, or 31. Ideally, an envelope is defined by fixing the cosine factor in the formula (eq. 1) at +1 or -1, so the phase effect disappears.

$$U(n) = \max\big(C(n)\big)_{0\le \varphi<2\pi} = \max\left(C0 + A\cos\left(\frac{60.5-n}{10.3} * 2\pi - \frac{2}{3}\pi - \varphi\right)\right)_{0\le \varphi<2\pi} = C0 + A \qquad (2)$$

Here, $U(n)$ represents the upper envelope of $C(n)$. According to (eq. 2), C26, C29, and C31 should share the same upper envelope, since $U(n) = C0 + A$ is independent of $n$. However, the upper envelopes derived from the three cyclizabilities differ from each other (Supplementary Fig. 5a, middle). The assumption that $A$ is independent of $n$, which was first adopted in (eq. 1) and led to the definition of envelopes in (eq. 2), does not agree well with the data. Therefore, $A$ should be considered a variable dependent on the biotin position, or $A(n)$.

$$U(n) = C0 + A(n), \qquad where\ n = 26, 29, 31$$

Similarly, the lower envelopes would follow $L(n) = C0 - A(n)$. Also, the original formula fitting C0 is revised.

$$C(n) = C0 + A(n)\cos\left(\frac{60.5-n}{10.3} * 2\pi - \frac{2}{3}\pi - \varphi\right), \qquad where\ n = 26, 29, 31 \qquad (3)$$

The dependence of $A(n)$ on $n$ originates from the adapter sequence. The adapter sequence has an intrinsic tendency to bend in certain directions. Changing the biotin tether position, which affects the orientation of the favored bending direction of an adapter sequence, will systematically alter the cyclizability measurement for each $n$. Such rotational phasing effect of adapter sequence was removed by a trigonometric formula similar to (eq. 1). In this way, we minimized the dependence of $A(n)$ on $n$ and found the envelopes independent of $n$. Accordingly, C26, C29, and C31 were adjusted to $C26_{corr}$, $C29_{corr}$, and $C31_{corr}$ in order that they share the same corrected envelopes that are independent of $n$. Then, $C0_{corr}$ was obtained by the average of the corrected upper and lower envelopes at each position. $C0_{corr}$ obtained in this way is minimally affected by the adapter sequences, and the strand in which the DNA sequence is read is independent of $C0_{corr}$ (Fig. 1f).

**Defining adapter-corrected cyclizability**

Envelope was separated into a constant term and a trigonometric term.

$$U(n) = U0 + A'\cos\left(\frac{60.5-n}{10.3} * 2\pi - \frac{2}{3}\pi - \varphi'\right)$$

$$L(n) = L0 + A''\cos\left(\frac{60.5-n}{10.3} * 2\pi - \frac{2}{3}\pi - \varphi''\right)$$

The trigonometric terms account for the rotational phasing of the adapter sequences. Approximation of $A'$ and $\varphi'$, or $A''$ and $\varphi''$ was done using the Python SciPy 1.9.3 package with the starting estimate of [1, 1] (PMID: 32015543).

The adapter-corrected cyclizability was defined as follows.

$$C(n)_{corr} := L0 + \left(C(n) - L(n)\right) * \frac{U0 - L0}{U(n) - L(n)}, \qquad where\ n = 26, 29, 31$$

The rotational phases of the 50 bp variable sequences create oscillatory modulations in $C(n)_{corr}$ that are bounded by $L0$ and $U0$ (Supplementary Fig. 5b). We defined $C0_{corr}$ as an average of $L0$ and $U0$ (Supplementary Fig. 5a, right, Supplementary Fig. 5b).

$$C0_{corr} := \frac{U0 + L0}{2}$$

Notably, $C0_{corr}$, is a value independent of the strand (original or reverse complementary) of an input sequence (Fig. 1f). This confirms that the effect of adapter sequences on cyclizability is minimized as expected.

Another option is to define $C0_{corr}$ as a weighted average of $C26_{corr}$, $C29_{corr}$, and $C31_{corr}$ following the earlier approach[1]. The method however conveys inaccuracy, as the choice of base pairs for a single helical turn (in this study, 10.3 bp) may vary depending on the sequence-specific structures of DNA. $C0_{corr}$ defined in this way shows a suboptimal correlation between the original and reverse complementary sequences (Fig. S3).

The A/T content shows an oscillatory pattern along 50 bp DNA sequences with the 1,000 highest or the 1,000 lowest $C26_{corr}$ in the Random library[1]. We reproduced similar results using $C29_{corr}$ or $C31_{corr}$ (Supplementary Fig. 5c). These results imply that $C26_{corr}$, $C29_{corr}$, and $C31_{corr}$ are the propensity of DNA looping towards three different directions in space, and A/T motifs in a specific position increase the bending of DNA molecule to a certain direction. On the other hand, $C0_{corr}$ is independent of the looping geometry of DNA, and A/T content is uniform across 50 bp DNA sequences with the 1,000 highest or the 1,000 lowest $C0_{corr}$ in the Random library (Supplementary Fig. 5c).

**Computational details**

The envelopes of C26 were determined by finding a curve that encloses the oscillatory pattern of C26 along a sufficiently long DNA sequence (longer than 100 bp to see clear oscillations). For an upper envelope, we collected the local maxima of C26 along the DNA sequence. A C26 value is a local maximum if it is the highest among 7 consecutive C26 values (including itself and three C26 values to upstream and downstream with a stride of 1 bp). We then outlined the set of local maxima using cubic interpolation by adopting the interp1d function of Python SciPy 1.9.3 package[2]. Occasionally, C26 is higher than the interpolated upper envelope. In such cases, we updated the upper envelope at the position with the C26 value. The lower envelope was determined in a similar way by interpolating the local minima instead.

We generated two DNA libraries of 150 bp sequences by appending random 50 bp sequences to both ends of the sequences of the Random library. Accordingly, the two libraries share the same 50 bp sequences in the middle, whose adapter-corrected cyclizability values were computed. The Pearson's $R$ between replicated calculations of adapter-corrected cyclizability is higher than 0.97, confirming that adapter-corrected cyclizability is a well-defined quantity independent of its context (Supplementary Fig. 5d).

$C0_{corr}$ correlates well with $C0_{corr}$ of a reverse complementary DNA (Pearson's $R$ = 0.966, Fig. 1e). Previous methods were designed to increase the correlation between C0 values of an original and the reverse complementary sequences by using information from both the forward and reverse strands as inputs (PMID: 35288750, 36866046, 37697433), which is unnecessary for training $C0_{corr}$.

In a previous study, the intrinsic cyclizability of 50 bp and 55 bp variable sequences was compared to confirm that the difference in rotational phase between ends is not a significant factor determining cyclizability. In this study, we trained the models with an input size of 55 bp (Fig. S4) directly on the loop-seq measured cyclizability of 12,288 55 bp variable sequences included in the library L. Due to the small size of library L, we omitted the step that screens sequences by uncertainty scores. Other parameters were unchanged. 12,472 55 bp sequences in library L share 50 bp from the left with sequences in the Random library. Using the cyclizability prediction model on 55 bp, $C0_{corr}$ of 55 bp is compared to $C0_{corr}$ of 50 bp from the left. The two adapter-corrected cyclizability correlates well despite the difference in information at the 3' end over 5 bp and the omission of a data refinement step in Library L (Pearson's $R$ = 0.863, Supplementary Fig. 5f).

#### Supplementary Note 5. Bending of $(dA)_n$ in 3-dimensional space

For each n from 0 to 20, we generated 500 random 101 bp DNA sequences with $(dA)_n$ in the middle surrounded by random sequences. We placed an equal number of bases to both ends to reach a total of 101 bp when n is odd, e.g., 5'-$N_{47}B(dA)_5BN_{47}$-3', whereas one more base was added to upstream of $(dA)_n$ when n is even, e.g., 5'-$N_{47}B(dA)_6BN_{46}$-3'. B implies non-dA bases (dT, dG, or dC).

For each length of poly(dA:dT) tract, the predicted $C26_{corr}$, $C29_{corr}$, and $C31_{corr}$ were represented by 500×52 matrices. Averaged $C26_{corr}$, $C29_{corr}$, and $C31_{corr}$ at each position (52-dimensional vectors) were used in spatial analysis.

6 and 17 bp poly(dA:dT) tracts have the biggest curving amplitudes in 3-dimensional space, whereas the 12 bp poly(dA:dT) tract has the smallest curving amplitude (Fig. S5a). This implies that a 6 bp poly(dA:dT) tract bends strongly in a single direction, and a 12 bp poly(dA:dT) tract has a rod-like structure with little preference for bending in a specific direction, as two adjacent 6 bp poly(dA:dT) tracts would bend in opposite directions. Spatial analysis in 3-dimensional space reveals an intrinsic looping tendency of 6 bp poly(dA:dT) tract towards the minor groove (Fig. S5b), as previously suggested[3].

We further verified the idea by varying the distance between two poly(dA:dT) tracts. To analyze the synergic effect between two poly(dA:dT) tracts on $C0_{corr}$, we generated 50 bp sequences containing two poly(dA:dT) tracts. For poly(dA:dT) tracts, we used $(dA)_n$ or $(dT)_n$, allowing four combinations. We placed two 6 bp poly(dA:dT) tracts separated by n bp, where n is ranging from 0 to 20. If n is odd, we placed one base on the upstream of the first poly(dA:dT) tract, e.g., 5'-$N_{16}B(dA)_6BN_4V(dT)_6VN_{15}$-3', and if n is even, we placed same length of random DNA upstream of the first and the downstream of the second poly(dA:dT) tract, e.g., 5'-$N_{15}B(dA)_6BN_4V(dT)_6VN_{15}$-3'. B and V imply non-dA and non-dT bases. For each n, we generated 500 random 50 bp sequences and predicted $C0_{corr}$ (Fig. S5c). We applied the same rule for placing 6 and 12 bp poly(dA:dT) tracts within a 50 bp DNA (Fig. S5c). Distance between two poly(dA:dT) tracts was a significant factor influencing $C0_{corr}$, especially when both tracts were 6 bp, reflecting the anisotropic bending of 6 bp poly(dA:dT) tracts (Fig. S5c).

#### Supplementary Note 6. Quantifying mechanical selection

#### Variables

In this section, we describe a detailed process for quantifying mechanical selection (Methods). When aligning the genomes of 1,011 yeast isolates to a 50 bp window, $C0_{corr}$ of the aligned natural sequences can be grouped into a few categories because natural genomes share common ancestors. The variables can be defined accordingly if there are $n$ distinct $C0_{corr}$ ($i$: integer ranging from 1 to $n$).

$c_i$: Counts of natural strains for each $C0_{corr}$.

$N$: Counts of all simulated $C0_{corr}$.

$r_i$: Rank of each natural $C0_{corr}$ among simulated $C0_{corr}$. Rank of 0 implies the lowest $C0_{corr}$ value.

$S \coloneqq \sum c_i r_i$, (or $\hat{S} \coloneqq \sum c_i \hat{r_i}$ when the ranks are determined).

We assumed $c_i$ is a constant and $r_i$ follow a uniform distribution, or, $r_i \sim U(0, N)$.

#### *Z*-score of mechanical selection

We calculated the expectation value and variance of $S$ as follows, assuming that $r_i$ are independent of each other.

$$E[S] = E[\sum c_i r_i] = E[r_i] \sum c_i = \frac{N \sum c_i}{2} \quad (1)$$

$$V[S] = \sum V[c_i r_i] = V[r_i] \sum c_i^2 = \frac{N(N+2) \sum c_i^2}{12} \quad (2)$$

Accordingly, the *Z*-score that describes mechanical selection is defined as follows.

$$Z = \frac{\hat{S} - E[S]}{(V[S])^{\frac{1}{2}}} \quad (3)$$

The computational complexity of the *Z*-score is linearly proportional to $n$, which does not exceed 100 in most cases.

#### *P*-value of mechanical selection

Under the null hypothesis, the two-tailed $P$-value is $2P(S \le \hat{S})$ when $\hat{S} \le E[S]$, or $2P(S \ge \hat{S})$ when $\hat{S} \ge E[S]$. For brevity, we will assume $\hat{S} \le E[S]$ from here, and the case with the opposite condition can be computed in a similar way.

One way of calculating the $P$-value is to count every possible combination of $r_i$ that satisfies $S \le \hat{S}$, say $T$, and obtain $2P(S \le \hat{S}) = 2T/(N+1)^n$, which requires a computational complexity proportional to $(N+1)^n$ to compute the exact value of $T$. To reduce computational complexity, we used two approaches to obtain an approximate $P$-value, as described in the following sections.

**Obtaining *P*-value by continuous approximation of discrete variables**

In this section, we assume that $r_i$ is a continuous uniform variable, producing similar results except in cases with very low $P$-values. In these cases, we use the dynamic programming approach described in the next section, which is based on discrete $r_i$ settings.

To simplify notations, we define $N_i := c_i N/2$, $X_i \sim U(-N_i, N_i)$, where $X_i$ is a continuous variable. In this case, $\widehat{X_i} = c_i \widehat{r_i}$ holds. Further, we define $X := S - E[S] = \sum X_i$. Similarly, $\hat{X} := \hat{S} - E[S] = \sum \widehat{X_i}$ holds. In this notation $E[X] = 0$ and $V[X] = V[S]$ hold.

We combine the known characteristic functions of $X_i$ (the Fourier transform of the probability density function), $\varphi_{X_i}(t) = E[e^{itX_i}]$, to infer the characteristic function of $X$, $\varphi_X(t)$, and obtain the probability distribution of $X$ using an inverse Fourier transform.

$$\varphi_X(t) = \varphi_{X_i}(t) \dots \varphi_{X_i}(t)$$

The following identity holds,

$$\varphi_{X_i}(t) := E[e^{itX_i}] = \int_{-N_i}^{N_i} \frac{e^{itx}}{2N_i} dx = \frac{\sin N_i t}{N_i t} = sinc(N_i t)$$

where

$$sinc(x) := \begin{cases} 0, & x = 0 \\ \dfrac{\sin x}{x}, & x \neq 0 \end{cases}$$

The characteristic function of $X$ is obtained as follows.

$$\varphi_X(t) = \varphi_{X_1}(t) \dots \varphi_{X_n}(t) = sinc(N_1 t) \dots sinc(N_n t) \tag{4}$$

Using the inversion formula[4], we obtain the probability distribution of $X$ from the characteristic function, leading to the cumulative probability distribution of $S$.

$$\begin{aligned} P(S < \hat{S}) &= P(X < \hat{X}) \\ &= \frac{1}{2} - \frac{1}{\pi}\int_0^\infty \frac{Im[e^{-itX}\varphi_X(t)]}{t}\, dt \\ &= \frac{1}{2} - \frac{1}{\pi}\int_0^\infty \frac{1}{t} sinc(-tX) sinc(N_1 t) \dots sinc(N_n t)\, dt \\ &= \frac{1}{2} + \frac{X}{\pi}\int_0^\infty sinc(tX) sinc(N_1 t) \dots sinc(N_n t)\, dt \\ &= \frac{1}{2} + \frac{\alpha X}{\pi}\int_0^\infty sinc(\alpha\omega X) sinc(\alpha N_1 \omega) \dots sinc(\alpha N_n \omega)\, d\omega \end{aligned} \tag{5}$$

where $t := \alpha\omega$ and $\alpha := \pi/(N_1 + \cdots + N_n)$.

The integrals can be replaced with summations[5].

$$P(S \leq \hat{S}) = P(X < \hat{X}) = \frac{1}{2} + \frac{\alpha X}{\pi}\sum_{t=1}^{\infty} sinc(\alpha X t) sinc(\alpha N_1 t) \dots sinc(\alpha N_n t) \tag{6}$$

The $sinc(\alpha X t) sinc(\alpha N_1 t) \dots sinc(\alpha N_n t)$ term converges to 0 quickly as $t$ increases, thus, the first few terms of the summation are sufficient to calculate a $P$-value with practical accuracy.

Errors after the first $p$ terms in the series (eq. 6) are calculated as follows.

$$e(p) = \left| P(S \leq \hat{S}) - \frac{1}{2} - \frac{\alpha X}{\pi} \sum_{t=1}^{p} sinc(\alpha X t) sinc(\alpha N_1 t) \dots sinc(\alpha N_n t) \right|$$

$$= \left| \frac{\alpha X}{\pi} \sum_{t=p+1}^{\infty} sinc(\alpha X t) sinc(\alpha N_1 t) \dots sinc(\alpha N_n t) \right|$$

$$\leq \frac{\alpha X}{\pi} \sum_{t=p+1}^{\infty} |sinc(\alpha X t) sinc(\alpha N_1 t) \dots sinc(\alpha N_n t)|$$

$$\leq \frac{\alpha X}{\pi} \sum_{t=p+1}^{\infty} \frac{1}{\alpha X t} \frac{1}{\alpha N_1 t} \cdots \frac{1}{\alpha N_n t} = \frac{1}{\pi N_1 \dots N_n \alpha^n} \sum_{t=p+1}^{\infty} \frac{1}{t^{n+1}}$$

$$\leq \frac{1}{\pi N_1 \dots N_n \alpha^n} \int_p^{\infty} \frac{1}{t^{n+1}} \, dt = \frac{1}{\pi N_1 \dots N_n \alpha^n p^n n} \tag{7}$$

We calculated the series (eq. 6) up to $p$ that makes the error $e(p)$ less than 1/1000 of the computed series. The series was extended until $p$ that satisfies the inequality below.

$$\left( \frac{1}{\pi N_1 \dots N_n \alpha^n p^n n} \right) \left( \frac{1}{\frac{1}{2} + \frac{\alpha X}{\pi} \sum_{t=1}^{p} sinc(\alpha X t) sinc(\alpha N_1 t) \dots sinc(\alpha N_n t)} \right) < \frac{1}{1000} \tag{8}$$

This error analysis is applicable only when $\hat{S} \leq E[S]$, as we initially assumed.

For $\hat{S} \geq E[S]$, we use inversed ranks, $N - r_i$, to have $\hat{S}' \leq E[S']$, and calculated $P(S' \leq \hat{S}')$ using the series summation until the error converges sufficiently as in (eq. 8). Then we get the *P*-value as below.

$$2P(S \geq \hat{S}) = 2P(S' \leq \hat{S}') \tag{9}$$

In case where a *P*-value is lower than $10^{-13}$ (which is smaller than the precision available in NumPy 1.24 and Numba 0.57 [6,7]), we used the *P*-value obtained through the dynamic programming described in the next section, as it provides a more reliable calculation for small *P*-values.

**Obtaining *P*-value by dynamic programming**

We used an algorithm based on dynamic programming to count all possible combinations of $r_i$ and return the $P$-value on a log-scale with base 10. The 2-dimensional array, dp[a, b], contains the number of possible combinations of $r_i$ $(1 \leq i \leq a)$, with $r_i = 0$ $(a < i \leq n)$ that produce the sum $S = b$. The algorithm has a complexity proportional to $n(\hat{S} + 1)$.

**Algorithm: Dynamic programming**

```
function pvalue_dp(cᵢ, Shat, N):
    n = length(cᵢ)
    dp = initialize 2D array of zeros with dimensions (n+1, Shat+1)
    dp[0][0] = 1

    M = 1
    cnt = 0

    for i from 1 to n:
        for j from 0 to Shat:
            idx = j - ((minimum(N, j//cᵢ[i-1]) + 1) * cᵢ[i-1])

            if idx >= 0:
                dp[i][j] += sum(dp[i-1][j:idx:-cᵢ[i-1]])
            else:
                dp[i][j] += sum(dp[i-1][j::-cᵢ[i-1]])

            if dp[i][j] > M:
                M = dp[i][j]

        if M > N + 1:
            M /= (N + 1)
            dp /= (N + 1)
            cnt += 1

    total_count = sum(dp[n])
    result = max(-log10(total_count) + log10(N+1) * (n-cnt) - log10(2), 0)
    return result # log10 of two-tailed P-value
```

**Information entropy indicates the diversity of natural variants**

We used information entropy to quantify the diversity of aligned natural genomic sequences. For a 50 bp window, there are $M$ alignments consisting of $k$ unique sequences, each containing $m_i$ $(i = 1, \ldots, k)$ strains $(M = \sum m_i)$. The information entropy, $H$, is defined as follows.

$$H \coloneqq -\left(\frac{m_i}{M}\right)\sum_{i=1}^{k}\log\left(\frac{m_i}{M}\right)$$

When the diversity of the aligned natural genomes is insufficient, *P*-values and *Z*-scores are not well reproduced in repeated calculations due to the randomness in generating the simulated mutations. Thus, a 50 bp window with an information entropy of aligned natural sequences lower than 0.75 was not considered in further analyses. The reproducibility is described in the following sections.

**Example: transcription start sites (TSS) of yeast population genomes**

The nucleosome-depleted region (NDR) upstream of the transcription start site (TSS) is a rigid DNA element, whose rigidity depends on gene expression levels. Highly expressed genes have a more rigid NDR[8]. Using the TSS DNA of yeast, we checked if the *P*-values and the *Z*-scores are reproducible.

TSS and their measured expression levels were determined using a previously described method[8]. We considered the yeast TSSs of all genes with both ends mapped with high confidence[9]. Expression levels of TSS were determined by the average RNA polymerase counts along the entire transcribed regions, or the first 500 bp in case the transcript is longer than 500 bp. The polymerase counts were obtained from NET-seq measurements[10]. The coordinates of TSS were mapped onto the sacCer3 assembly[11] using LiftOver[12]. The genomes of 1,011 yeast isolates[13] were aligned to the TSSs (-300 to 100 bp from TSS) of 6,420 genes.

**Reproducibility of *Z*-score and *P*-value**

The reproducibility of *Z*-score was evaluated by calculating the correlation between repeated *Z*-score computations for the same alignment of population genomes across 6,420 TSS regions. Correlation was below 0.9 across all 50 bp windows but increased up to 0.97 when selecting windows with an information entropy higher than 0.75. The correlation changed minimally with higher entropy thresholds (Fig. S6b). A similar pattern was observed when using *P*-value instead of *Z*-score (Fig. S6b).

We also checked if the non-uniform mutation frequency of each of the 6 possible nucleotide changes (dA>dT, dC>dT, etc.) could affect the calculation of *Z*-score and *P*-value. *Z*-score and *P*-value were calculated similarly in 6,420 TSS regions, but using previously reported relative mutation frequencies instead[14]. The corrected mutation frequency minimally affected the reproducibility of *Z*-score and *P*-value (Fig. S6c, d). In this study, we adopted uniform mutation rates for all possible nucleotide changes.

**On the false discovery rate**

To verify that our method accurately reports mechanical selection, we examined the distribution of *Z*-scores and *P*-values of simulated random variants generated under no selection pressure across 6,420 TSS regions. The simulated random variants are generated from natural alignments by preserving the number of unique sequences, the counts of each unique sequence, and the Hamming distance. If there are 500 yeast isolates with identical 50 bp DNA containing 3 mutations, we generate 500 identical 50 bp DNA sequences, each with 3 random mutations, in the pool of hypothetical alignments. Variants are generated randomly according to the relative frequency of 6 possible nucleotide changes[14]. *Z*-score was near 0 for the simulated variants under no selection pressure, as expected (Fig. S7a).

Quantile-quantile (Q-Q) plots between expected and predicted distribution of *P*-values can reveal whether our method reports excessive false discovery. When including all 50 bp windows of 6,420 TSS regions, Q-Q plot deviates significantly from the 1:1 line (the identity line) under no selection pressure, which serves as evidence of false discovery. The Q-Q plot converges to the 1:1 line when we select 50 bp windows with entropy higher than 0.75, supporting the idea that analyzing 50 bp windows with entropy higher than a threshold can reduce the false discovery rate (Fig. S7b).

In contrast, the Q-Q plot using *P*-values obtained from the natural alignments deviates from the 1:1 line after we selected regions with entropy higher than 0.75, which implies true mechanical selection (Fig. S7c). From the distribution of *P*-values (calculated from the natural alignments) in regions with entropy higher than 0.75, we obtained a false discovery rate of 0.2041 at a significance level of $P$-value $< 0.01$ (Fig. S7d).

**Interpreting and visualizing the results**

When plotting cyclizability (or *Z*-score), we labeled the x-axis with the base in the center of 50 bp windows (e.g. the x-label for a *Z*-score of 1 to 50 bp is 25). 50 bp windows under mechanical selection were highlighted in red or blue, indicating anti- or pro-rigidity selection, respectively. *Z*-score, *P*-value, and entropy are not plotted in most figures for brevity (Fig. S8).

**Supplementary Note 7: Details for quantifying mechanical selection in centromeres**

We defined 48 DNA templates, each 180 bp in length, that cover 16 centromeres. Three DNA templates with strides of 90 bp cover one centromere. When a yeast centromere is even base pairs-long, (length/2 – 90) bp from the upstream end and (90 – length/2) bp from the downstream end are selected as the 5' and 3' ends of the template in the middle among the three templates covering one centromere. When the length of centromere is odd, (length/2 – 89.5) bp from the upstream end and (90.5 – length/2) bp from the downstream end are selected as the 5' and 3' ends of the middle template. The upstream and downstream templates are defined by shifting the position of the middle template 90 bp upstream or downstream, respectively.

Blat[15] was used to align genomic sequences of 1,011 yeast strains[13]. Alignments with over 80% sequence match without indel mutations were selected. There were no regions with the number of alignments higher than 1,011*1.2, or zero alignments. *Z*-score and *P*-value of each 50 bp window were determined, and the results from three covers (upstream, middle, and downstream covers for each centromere) were merged as follows.

The following windows were concatenated:

The 1$^{st}$ to 111$^{th}$ 50 bp windows from the upstream cover,

The 22$^{th}$ to 110$^{th}$ 50 bp windows from the middle cover,

The 21$^{th}$ to 131$^{th}$ 50 bp windows from the downstream cover.

**Supplementary Note 8: Heuristics to accelerate cyclizability predictions for long DNA sequences**

Predicting cyclizability at Gbps length scale (e.g., a chromosome) by sliding 50 bp DNA windows with a stride of 1 bp has a computational complexity linearly proportional to the length of the DNA (Fig. S9a). This may increase the computational time and limit practical usage, especially at a population genome scale. However, any two neighboring cyclizability predictions are derived from two 50 bp DNA sequences that contain an overlapping 49 bp region, in which computations are repeated. In this section, we introduce heuristic methods to reduce this redundancy and shorten the computational time.

The first method is to use non-overlapping 50 bp windows, where the DNA segment is quantized into discrete 50 bp sequences, which are then processed sequentially (Fig. S9b). While the complexity of this algorithm is still linearly dependent on the length of the input BP, performing an order of magnitude fewer calculations results in a significantly faster processing time. To compare the output of the quantization method to the original sliding window method, we compared the averaged $C0_{corr}$ value for every computed window on randomly generated sequences. To do this, 100 random sequences of 1,000, 10,000, and 100,000 bp were generated and the sliding window and quantization algorithms were applied. Each calculated output was then summed together to yield an average $C0_{corr}$ value for each of the long sequences for comparison (for a 1,000 bp DNA as an example, the average of the 951 and 20 values of $C0_{corr}$ predicted using the sliding window and quantization algorithms, respectively). Pearson's $R$ between these two algorithms was ~ 0.8 (Fig. S9c). It is worthwhile to note here that this algorithm no longer provides the cyclizabilty at any given nucleotide (influenced by the surrounding bases) but provides a single value for each non-overlapping window. However, depending on the application, this greatly reduces computational run time.

The second method is to train a model that has an input of 1,000 bp DNA and an output of 951 cyclizability values per single prediction (Fig. S9d). The original model in this study contains two convolutional layers (Methods, Supplementary Fig. 3a). The first convolutional layer scans every 7-mer in a 50 bp DNA, and this computation is identical for an overlapping 49 bp region between two consecutive 50 bp windows. Thus, computing convolutional layers once for a long DNA and reusing these values multiple times is more efficient than computing convolutional layers repeatedly. We designed the model as follows (Fig. S9e). The model was implemented using Keras[16].

Input (200,) - A 1,000 bp DNA is converted into a 4,000-dimensional vector by one-hot encoding:

A: [1, 0, 0, 0], T: [0, 1, 0, 0], G: [0, 0, 1, 0], C: [0, 0, 0, 1]

First 1D convolution layer - Kernel size: 28, Output channels: 64, Stride: 4, Output shape: (994, 64). A bias term and a rectified linear unit (ReLU) activation were added.

Second 1D convolution layer - Kernel size: 33, Output channels: 32, Output shape: (962, 32). A bias term and a rectified linear unit (ReLU) activation were added.

Third 1D convolution layer - Kernel size: 12, Output channels: 50, Output shape: (951, 50). A bias term and a rectified linear unit (ReLU) activation were added.

Flatten layer - Output shape: (47550, ).

Fully connected layer (output) - Output shape: (951, ). This layer predicts 951 cyclizability values of a 1,000 bp DNA sequence.

30,000 random 1,000 bp DNA sequences were generated, and their $C0_{corr}$ (951 values per DNA sequence) were predicted using the original model (Methods, Supplementary Fig. 3a). This dataset was split into a 9:1 ratio for the training and testing datasets. The model predicting 1,000 bp at a time (Fig. S9d, e) was trained for 5 epochs, similar to the original model (Methods). Pearson's *R* between the predictions of the original and the newly trained model was 0.988 for the training dataset (Fig. S9f).

To compare the output of the model predicting 1,000 bp at a time to the original model, we compared the averaged $C0_{corr}$ value for every computed window on randomly generated sequences. 100 random sequences of 1,000, 10,000, and 100,000 bp were generated, and their $C0_{corr}$ values were calculated using the two methods. Each calculated output was then summed together to yield an average $C0_{corr}$ value for each of the long sequences. Pearson's *R* between the two methods was ~ 0.99 (Fig. S9g).

We compared the speed of cyclizability prediction for three different methods with varying DNA lengths on Google Colaboratory (Retrieved Nov. 6, 2024, https://colab.research.google.com/; Intel Xeon CPU with 2 vCPUs (virtual CPUs) and 13GB of RAM). The prediction time was proportional to the length of DNA in all three methods (Fig. S9h). For a 1,000,000 bp DNA, the sliding method (Fig. S9a) took 127.85 seconds, the non-overlapping windows method (Fig. S9b) took 2.13 seconds, and the 1 kbp prediction model (Fig. S9d, e) took 10.93 seconds (Fig. S9h).

**Fig. S1**

Uncertainty score calculated by frequentist statistics and the related 95% CI calculated by Bayesian statistics for the measured C26 of the Tiling library.

**Fig. S2**

Measured vs predicted cyclizability of the sequences in the Tiling library containing 7-mer motifs, 5'-CGAGAAG-3' or 5'-CTTCTCG-3'. Models were trained on the dataset that has no such 7-mer motifs (Supplementary Note 3).

**Fig. S3**

$C0_{corr}$ of original vs the related reverse complementary sequence, when using an alternative definition of $C0_{corr}$ in Supplementary Note 4.

**Fig. S4**

Structure of the CNN model with the modified input size of 55 bp.

**Fig. S5**

**a**, $C26_{corr}$ and the bending amplitude of 500 sequences of 101 bp, each containing 0 to 20 bp poly(dA:dT) tracts in the middle. **b**, Average bending propensity of DNA sequences containing a 6 bp poly(dA:dT) tract at the center. **c**, Average $C0_{corr}$ of 50 bp sequences containing two poly(dA:dT) tracts for various sized gaps between the tracts. Two 6 bp poly(dA:dT) tracts, or 6 and 12 bp poly(dA:dT) tracts, are separated by a gap. For each gap size, n=500. $C0_{corr}$ for 12,472 50 bp sequences chosen at random, or containing $(dA)_6$, or $(dT)_6$ in the middle was plotted separately. Whisker box plots are shown together with the scattered data.

**Fig. S6**

Pearson's correlation between repeated calculations of *Z*-scores, **a**, and *P*-values, **b**. **a**, **b**, For the analysis, 50 bp regions at 6,420 TSS of yeast were selected by entropy higher than certain threshold values as indicated in the figure. **c**, Correlation between the original *Z*-score and the *Z*-score calculated using a different mutation rate setting (PMID: 24847077), among 50 bp regions at 6,420 TSS with entropy higher than certain threshold values as indicated. **d**, Repeat of **c** for *P*-values. **a-d**, Shaded backgrounds denote 95% CI of Pearson's correlation.

**Fig. S7**

**a**, *Z*-score of yeast TSS (left), and *Z*-score of yeast TSS with simulated random mutations (right), grouped by expression levels. The simulated random variants preserve the Hamming distance and the multiplicity of TSS sequences. For example, if there were 500 yeast isolates sharing a TSS sequence with 5 natural mutations, then we generated the corresponding 500 simulated TSS sequences sharing 5 random mutations. Shaded backgrounds denote s.e.m. **b**, Expected vs observed distribution of *P*-value in 6,420 TSS regions that have simulated random variants. **c**, Expected vs observed distribution of *P*-value in 6,420 TSS regions with natural variants. **b**, **c**, For the analysis, 50 bp regions were selected by entropy higher than certain threshold values as indicated. **d**, Distribution of *P*-values in 6,420 TSS regions with natural mutations. 50 bp windows with entropy higher than 0.75 were used in the analysis.

**Fig. S8**

Visualizing mechanical selection in a genomic region. 50 bp windows with information entropy higher than 0.75 and *P*-value lower than 0.01 are considered mechanical selection.

**Fig. S9**

**a**, Schematic of 50 bp windows sliding across a long DNA sequence with a stride of 1 bp. **b**, Schematic of non-overlapping 50 bp windows covering a long DNA sequence. **c**, $C0_{corr}$ predicted using sliding windows vs non-overlapping windows averaged over each random DNA sequence. The analyses were done individually for DNA sequences of 1,000, 10,000, and 100,000 bp. Pearson's correlation and sample size are shown. **d**, Schematic of the accelerated prediction method. Cyclizability values for a 1,000 bp DNA sequence were computed by a single prediction. **e**, Model structure for predicting 951 cyclizability values for a 1,000 bp DNA sequence. **f**, $C0_{corr}$ predicted using the original model vs the model in **e** (Supplementary Note 8) for the training dataset. Pearson's correlation and sample size were shown. **g**, $C0_{corr}$ predicted using sliding windows vs the model computing 1,000 bp per run (Supplementary Note 8) averaged over each random DNA sequence. The analyses were done individually for DNA sequences of 1,000, 10,000, and 100,000 bp. Pearson's correlation and sample size are shown. **h**, Computation time to predict $C0_{corr}$ using three different methods (panel **a**, **b**, and **d**) at various DNA lengths. For each bar, n=3.

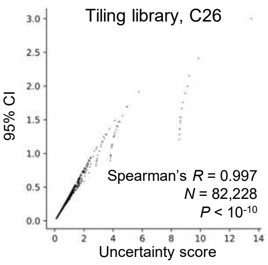
Tiling library, C26
95% CI
Spearman's R = 0.997
N = 82,228
P < 10-10
Uncertainty score
0.0
0.5
1.0
1.5
2.0
2.5
3.0
0
2
4
6
8
10
12
14

Fig. S1

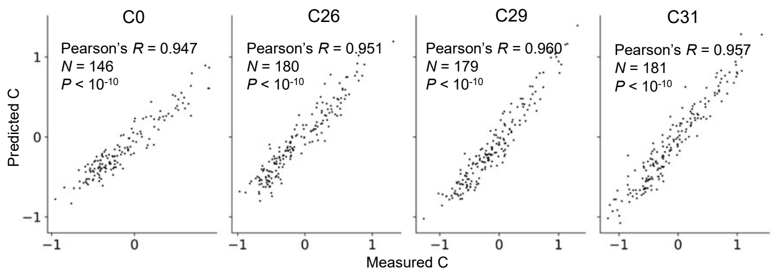
C0
Pearson's R = 0.947
N = 146
P < 10-10
C26
Pearson's R = 0.951
N = 180
P < 10-10
C29
Pearson's R = 0.960
N = 179
P < 10-10
C31
Pearson's R = 0.957
N = 181
P < 10-10
Predicted C
Measured C

Fig. S2

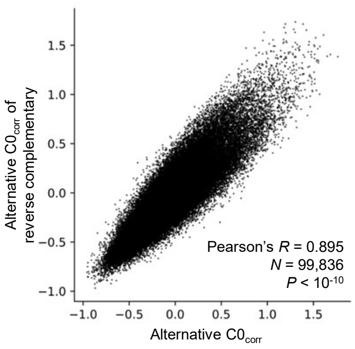
Alternative $C0_{corr}$ of reverse complementary
Alternative $C0_{corr}$
Pearson's $R$ = 0.895
$N$ = 99,836
$P < 10^{-10}$
1.5
1.0
0.5
0.0
−0.5
−1.0
−1.0
−0.5
0.0
0.5
1.0
1.5

Fig. S3

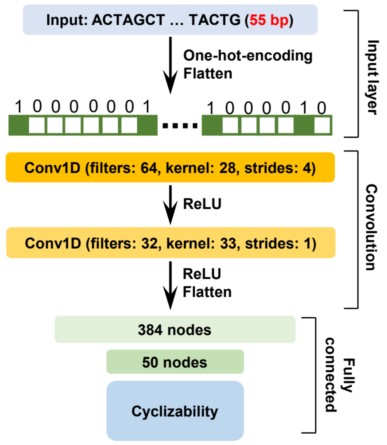
Input: ACTAGCT … TACTG (55 bp)
One-hot-encoding
Flatten
1 0 0 0 0 0 0 1
1 0 0 0 0 0 1 0
Input layer
Conv1D (filters: 64, kernel: 28, strides: 4)
ReLU
Conv1D (filters: 32, kernel: 33, strides: 1)
ReLU
Flatten
Convolution
384 nodes
50 nodes
Cyclizability
Fully connected

Fig. S4

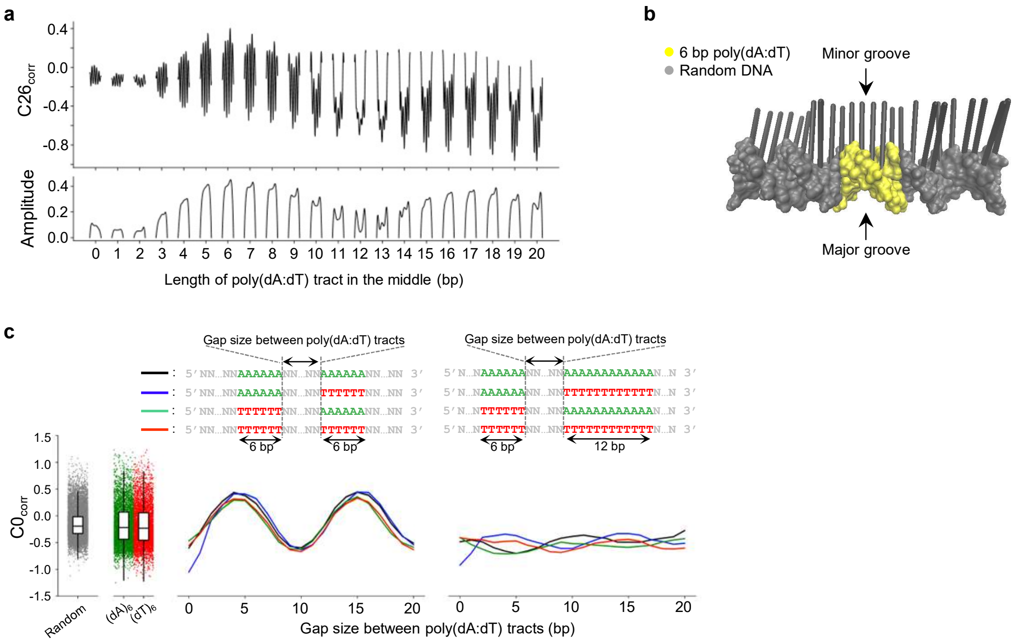
a
C26corr
Amplitude
Length of poly(dA:dT) tract in the middle (bp)
b
6 bp poly(dA:dT)
Random DNA
Minor groove
Major groove
c
Gap size between poly(dA:dT) tracts
6 bp
12 bp
C0corr
Random
(dA)6
(dT)6
Gap size between poly(dA:dT) tracts (bp)

Fig. S5

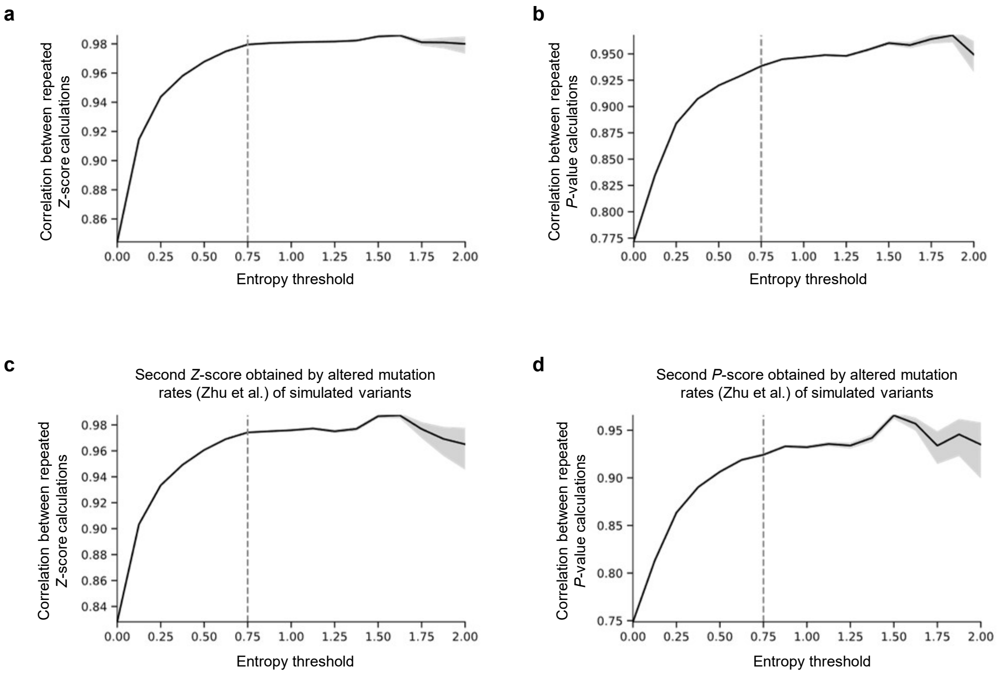
a
Correlation between repeated Z-score calculations
Entropy threshold
b
Correlation between repeated P-value calculations
Entropy threshold
c
Second Z-score obtained by altered mutation rates (Zhu et al.) of simulated variants
Correlation between repeated Z-score calculations
Entropy threshold
d
Second P-score obtained by altered mutation rates (Zhu et al.) of simulated variants
Correlation between repeated P-value calculations
Entropy threshold

Fig. S6

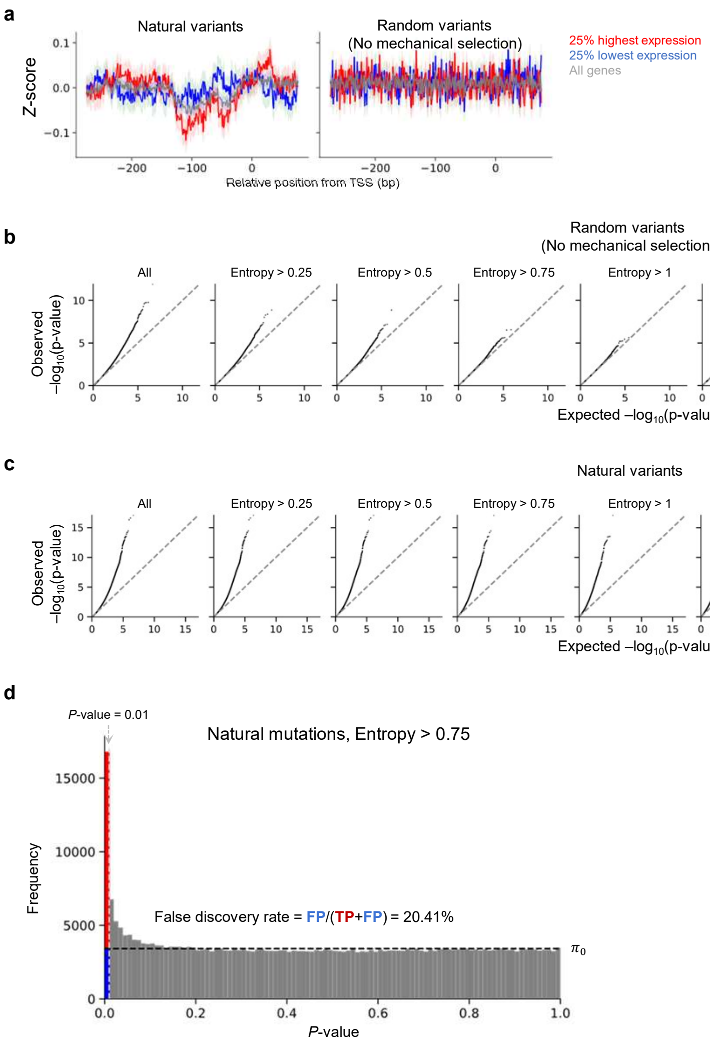

a
Natural variants
Random variants
(No mechanical selection)
25% highest expression
25% lowest expression
All genes
Z-score
Relative position from TSS (bp)
b
Random variants
(No mechanical selection)
All
Entropy > 0.25
Entropy > 0.5
Entropy > 0.75
Entropy > 1
Entropy > 1.25
Entropy > 1.5
Entropy > 1.75
Entropy > 2
Observed –log10(p-value)
Expected –log10(p-value)
c
Natural variants
All
Entropy > 0.25
Entropy > 0.5
Entropy > 0.75
Entropy > 1
Entropy > 1.25
Entropy > 1.5
Entropy > 1.75
Entropy > 2
Observed –log10(p-value)
Expected –log10(p-value)
d
P-value = 0.01
Natural mutations, Entropy > 0.75
Frequency
False discovery rate = FP/(TP+FP) = 20.41%
π0
P-value


Fig. S7

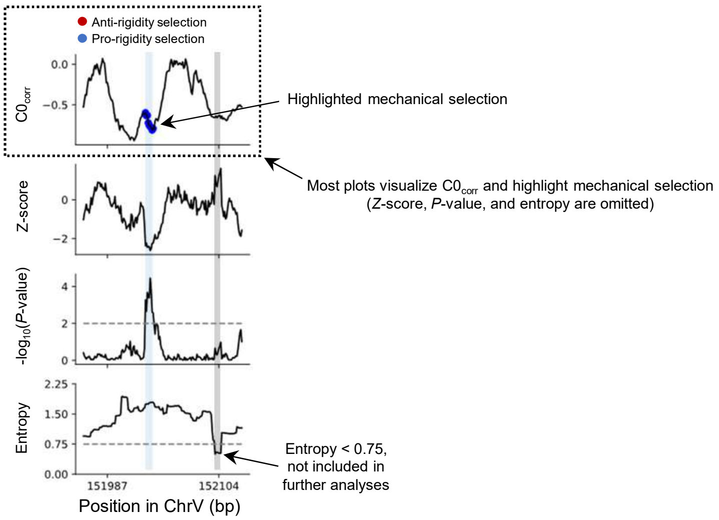

Anti-rigidity selection
Pro-rigidity selection
C0corr
0.0
−0.5
Highlighted mechanical selection
Most plots visualize C0corr and highlight mechanical selection
(Z-score, P-value, and entropy are omitted)
Z-score
0
−2
-log10(P-value)
4
2
0
Entropy
2.25
1.50
0.75
0.00
151987
152104
Position in ChrV (bp)
Entropy < 0.75,
not included in
further analyses


Fig. S8

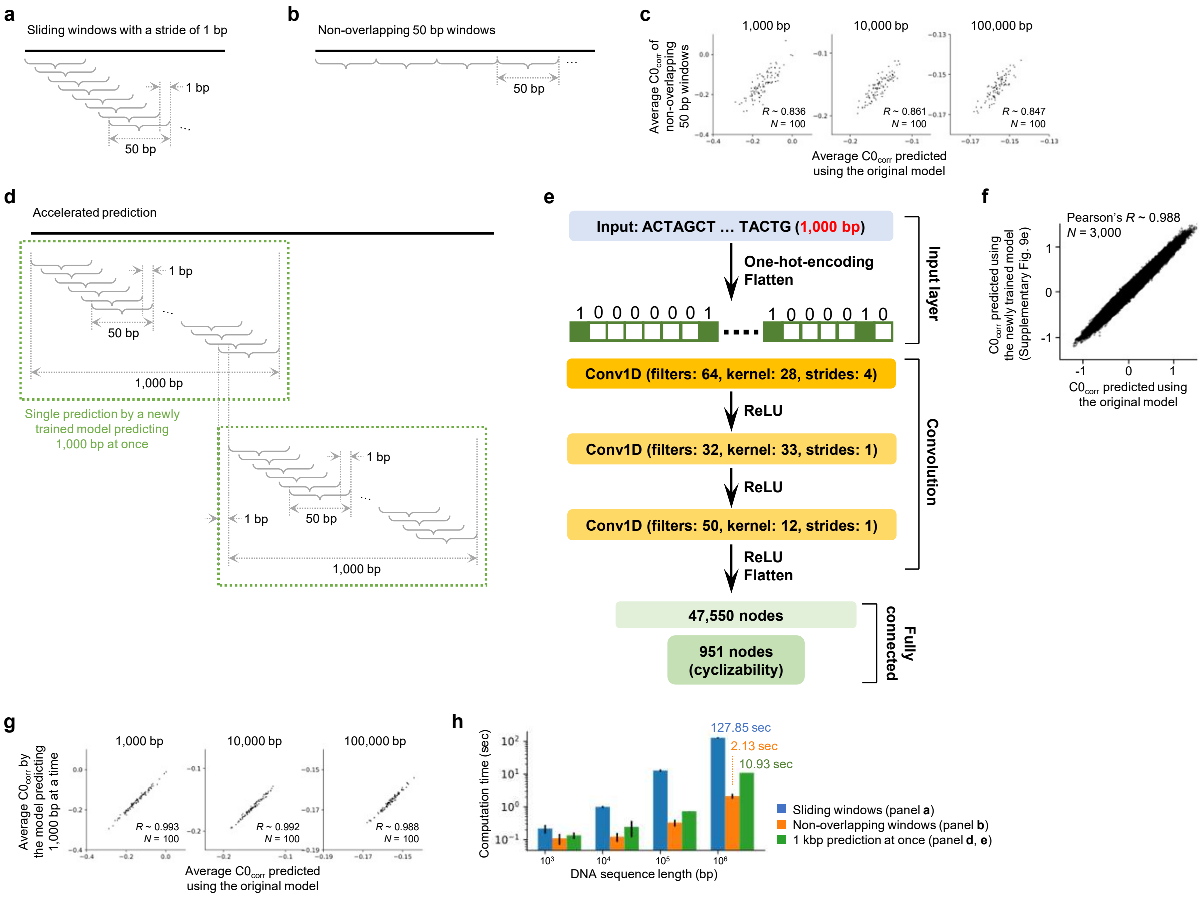
a
Sliding windows with a stride of 1 bp
1 bp
50 bp
b
Non-overlapping 50 bp windows
50 bp
c
1,000 bp
10,000 bp
100,000 bp
Average C0corr of non-overlapping 50 bp windows
R ~ 0.836
N = 100
R ~ 0.861
N = 100
R ~ 0.847
N = 100
Average C0corr predicted using the original model
d
Accelerated prediction
1 bp
50 bp
1,000 bp
Single prediction by a newly trained model predicting 1,000 bp at once
e
Input: ACTAGCT … TACTG (1,000 bp)
One-hot-encoding
Flatten
Input layer
Conv1D (filters: 64, kernel: 28, strides: 4)
ReLU
Conv1D (filters: 32, kernel: 33, strides: 1)
ReLU
Conv1D (filters: 50, kernel: 12, strides: 1)
ReLU
Flatten
Convolution
47,550 nodes
951 nodes (cyclizability)
Fully connected
f
Pearson's R ~ 0.988
N = 3,000
C0corr predicted using the newly trained model (Supplementary Fig. 9e)
C0corr predicted using the original model
g
1,000 bp
10,000 bp
100,000 bp
Average C0corr by the model predicting 1,000 bp at a time
R ~ 0.993
N = 100
R ~ 0.992
N = 100
R ~ 0.988
N = 100
Average C0corr predicted using the original model
h
127.85 sec
2.13 sec
10.93 sec
Computation time (sec)
DNA sequence length (bp)
Sliding windows (panel a)
Non-overlapping windows (panel b)
1 kbp prediction at once (panel d, e)

Fig. S9
